# Preventing neuropathy and improving anti-cancer chemotherapy with a carbazole-based compound

**DOI:** 10.1101/2025.03.10.642317

**Authors:** Lauriane Bosc, Maria Elena Pero, David Balayssac, Nathalie Jacquemot, Jordan Allard, Peggy Suzanne, Julien Vollaire, Cécile Cottet-Rousselle, Sophie Michallet, Joran Villaret, Sakina Torch, Sumari Marais, Bénédicte Elena-Herrmann, Uwe Schlattner, Anne Mercier, Véronique Josserand, Chantal Thibert, Patrick Dallemagne, Francesca Bartolini, Laurence Lafanechère

**Affiliations:** Institute for Advanced Biosciences, Université Grenoble Alpes, INSERM U1209, CNRS UMR 5309; Grenoble, France; Department of Pathology and Cell Biology, Vagelos College of Physicians and Surgeons, Columbia University; New York, NY 10032, USA; Department of Veterinary Medicine and Animal Production, University of Naples Federico II; 80137 Naples, Italy; Université Clermont Auvergne, U1107, NEURO-DOL, INSERM, CHU Clermont-Ferrand, Direction de la recherche clinique et de l’innovation; Clermont-Ferrand, France; Université Clermont Auvergne, U1107, NEURO-DOL, INSERM, Clermont-Ferrand, France; Université Normandie, UNICAEN, CERMN; 14032 Caen, France; Laboratory of Fundamental and Applied Bioenergetics, Université Grenoble Alpes, INSERM U1055; Grenoble, France; Department of Physiology, School of Medicine, Faculty of Health Sciences, University of Pretoria; Pretoria 0028, South Africa; Institut Universitaire de France (IUF); 75231 Paris, France

## Abstract

Advances in cancer treatment have led to a steady increase in the rate of disease remission. However, while many treatment-related adverse effects gradually resolve after therapy, chemotherapy-induced peripheral neuropathy (CIPN) often persists, with no means of prevention or direct treatment available. Herein, we present Carba1, a novel bi-functional carbazole that mitigates neuropathy through two distinct mechanisms. First, by interacting with tubulin, Carba1 reduces the required dose of taxanes, widely used chemotherapy drugs notorious for their toxic side effects, including CIPN. Second, Carba1 activates nicotinamide phosphoribosyltransferase (NAMPT), the rate-limiting enzyme in the NAD salvage pathway, triggering a metabolic rewiring that enhances the resilience of neurons and Schwann cells against chemotherapy-induced toxicity. We demonstrate the neuroprotective efficacy of Carba1 both *in vitro,* against neurotoxicity induced by paclitaxel (PTX), cisplatin, and bortezomib, and *in vivo* in a rat model of PTX-induced neuropathy. Importantly, we establish that Carba1 does not compromise the therapeutic efficacy of PTX nor promotes tumor growth. Comparative analyses of Carba1 derivatives further suggest the potential of designing compounds with either dual synergistic and neuroprotective activity or exclusive neuroprotective properties. Altogether, our findings position Carba1 as a promising therapeutic candidate for preventing CIPN, with the potential, if successfully translated to clinical settings, to improve both the quality of life and treatment outcome for cancer patients.

## INTRODUCTION

Microtubules (MTs) are dynamic cytoskeletal filaments with crucial roles in cell division and physiology, which depend on their rapid remodeling through polymerization and depolymerization. Targeting this finely tuned process is a major therapeutic strategy in oncology (for review, see (*1*)) and drugs that interfere with MTs are key components of chemotherapies for treating carcinomas. Among them, paclitaxel (PTX) stands out as one of the most successful chemotherapeutic agents (*2*). PTX binds to the taxane site tubulin and stabilizes the MT lattice by strengthening tubulin contacts (*3*). At stoichiometric concentrations, PTX promotes MT assembly whereas at low and clinically relevant concentrations, PTX primarily suppresses MT dynamics without significantly affecting MT-polymer mass (*4*). However, MT-interfering drugs can face resistance and cause adverse effects like neutropenia, gut toxicity, alopecia, and peripheral neuropathies (*4*, *5*).

Chemotherapy-induced peripheral neuropathy (CIPN) is a debilitating adverse effect of neurotoxic anticancer drugs, affecting millions of patients annually. Characterized by distal bilateral tingling, burning, numbness and neuropathic pain of the limbs (*6*), CIPN affects about two-thirds of patients treated with chemotherapy (*7*). CIPN can lead to reduced chemotherapy doses, thus compromising remission chances. Unlike other chemotherapy side effects, neuropathies often persist after remission in about 25% of survivors, representing a significant public health concern (*6*, *8*, *9*). Currently, there are no preventive strategies, and only duloxetine has shown moderate success in managing CIPN pain (*10*, *11*).

To address resistance and CIPN toxicity with PTX, we explored compounds that, when used alone, show no significant biological effects but can significantly enhance the efficacy of PTX, enabling a reduction in PTX dosage. We previously discovered that the compound Carba1, a derivative of the carbazole series, sensitizes cells to a low, non-toxic dose of PTX (*12*). Carba1 also showed synergy with PTX *in vivo* in xenografted mice. It binds with low affinity to tubulin’s colchicine site, placing MTs in a favorable state for PTX binding (*12*).

In this study, we demonstrate that Carba1 synergizes with other compounds targeting the tubulin taxane site, thereby enhancing our understanding of its structural mechanism of action on tubulin. We also provide evidence that Carba1 remarkably protects neurons from CIPN, both *in vitro* and in a rat model. We found that the neuroprotective action of Carba1 results from the stimulation of NAD biosynthesis through activation of the enzyme nicotinamide phosphoribosyltransferase (NAMPT), a neuroprotective mechanism applicable to various types of CIPN beyond those induced by taxanes (*13*, *14*). Finally, our structure-activity relationship analysis shows that the carbazole series can be chemically modulated to prioritize neuroprotection, offering a promising therapeutic strategy to prevent CIPN.

## RESULTS

### The synergistic effect of Carba1 is specific to compounds that bind to the taxane site on tubulin

We previously demonstrated that Carba1, with minimal cytotoxicy in dividing cells with a GI_50_ (50% of growth inhibition) higher than 25 µM after a 72h treatment, synergizes PTX (*12*). To determine whether this synergy extends to similar compounds, we analyzed the cytotoxicity induced by Carba1 in combination with docetaxel (DTX), nab-paclitaxel (nab-PTX, Abraxane®) and epothilone-B (Epo-B) on HeLa cells (Fig. 1A, B). DTX, which was originally extracted from the needles of the European yew Taxus baccata in the 1980s, differs from PTX, which was first extracted from the bark of the Pacific yew Taxus brevifolia in the 1960s. The difference lies in the 10’ position of the baccatin ring and the 3’ position of the PTX side ring. Nab-PTX is a more soluble albumin-conjugated and a recently marketed formulation of PTX, while Epo-B is a 16-membered macrolide which differs structurally from the taxanes but binds to tubulin at the taxane site (*3*). We also tested two drugs frequently used in cancer chemotherapy, the alkylating agent cisplatin (Cis) and the reversible inhibitor of the 26S subunit of the proteasome, bortezomib (Bort, Fig. 1A). While there is no evidence that Cis might affect tubulin, Bort has been shown to stabilize MTs in neurons (*15*, *16*) although the mechanism underlying this stabilization appears to be different from that of taxanes (*17*, *18*) and remains to be fully elucidated. As shown in Fig. 1B, we found that DTX (GI_50_ of 1.1 ± 0.2 nM) was more potent than PTX, which has a GI_50_ of 4.2 ± 0.4 nM, as previously described (*19*). On the other hand, nab-PTX was less cytotoxic, with a GI_50_ of 6.0 ± 0.9 nM. Epo-B was highly cytotoxic, with a GI_50_ of 0.5 ± 0.03 nM. The addition of 12 µM Carba1 significantly reduced the GI_50_ of all the tested drugs that impairs MTs dynamics (PTX, DTX, nab-PTX and Epo-B by 2.6-fold, 3.8-fold, 1.8-fold, and 4.4-fold, respectively) indicating that Carba1 exerts a synergistic effect with other compounds sharing the ability to bind to the taxane site on tubulin. In contrast, the GI_50_ of Cis and Bort (166.7 ± 30.8 nM and 35.9 ± 3.7 nM, respectively) were not significantly affected when these compounds were used in combination with Carba1 (235.8 ± 42.6 nM and 40.0 ± 3.3 nM, respectively), and instead a slight upward trend was observed (Fig. 1B). Taken together, these results indicate that the synergistic effect of Carba1 requires occupation of the tubulin taxane site, supporting our earlier findings on Carba1’s synergistic mechanism, which involves alterations in MT-lattice regions at the growing end, promoting binding of compounds to the taxane site (*12*).

**Fig. 1.**
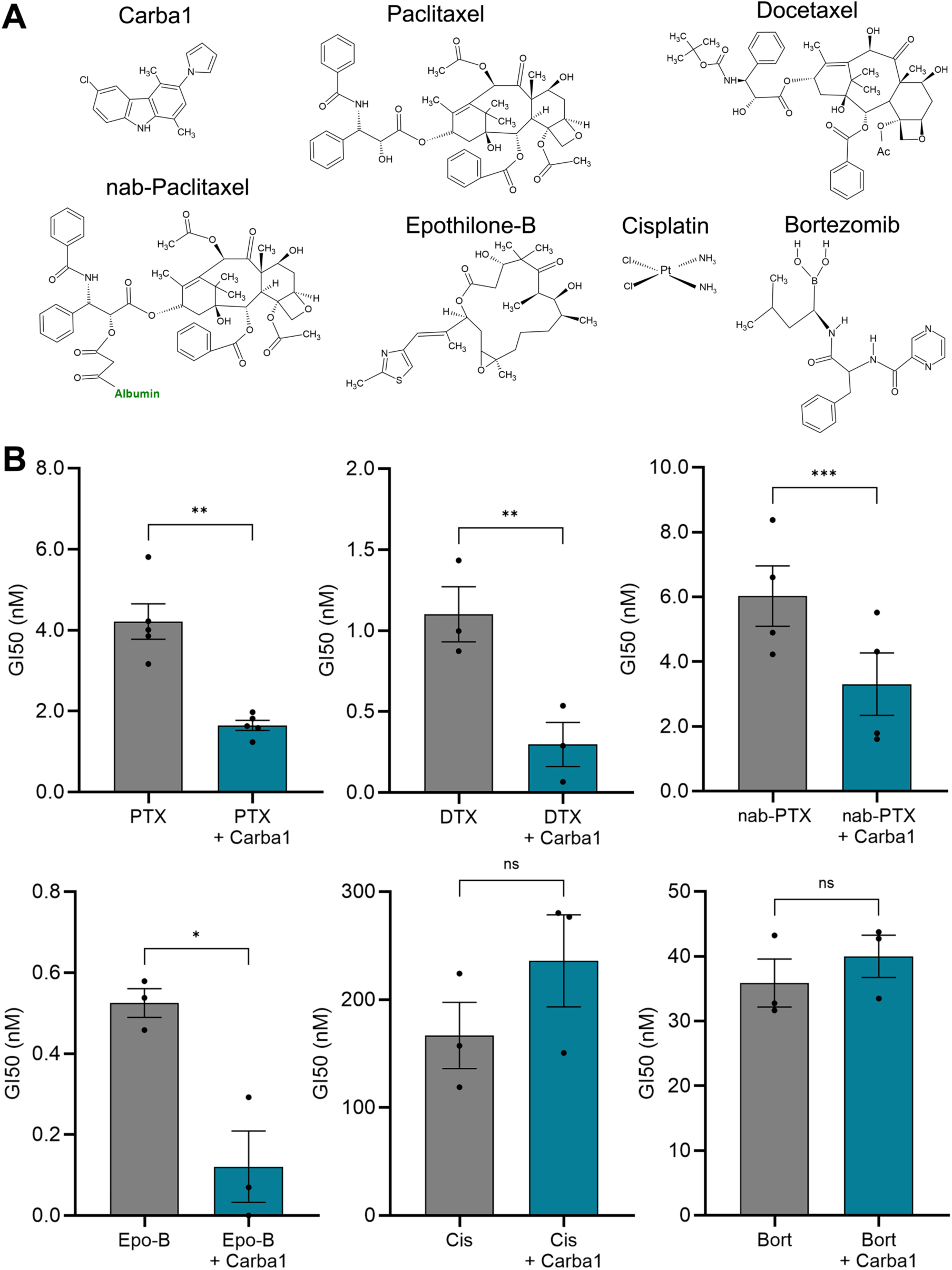
Carba1 synergizes only with chemotherapeutic agents that bind to the taxane site on tubulin. **(A)** Chemical structures of the compounds tested. **(B)** Effects of selected chemotherapeutic agents – Paclitaxel (PTX), Docetaxel (DTX), nab-Paclitaxel (nab-PTX), Epothilone-B (Epo-B), Cisplatin (Cis), Bortezomib (Bort) - on viability of HeLa cells alone or in combination with Carba1. Cells were incubated for 72 h with the indicated concentrations of the drugs with or without 12 µM of Carba1. The percentage of viable cells was calculated following a Prestoblue assay and shown as GI_50_ (50% of growth inhibition) of the drug when applied alone (grey) or in combination with Carba1 (blue). Data are presented as mean ± SEM of at least 3 independent experiments.* p< 0.05, ** p<0.005, ***p<0.0005, ns non-significant, t-Test.

### Carba1 prevents PTX-associated CIPN *in vitro* and *in vivo*

We previously reported that the combined administration of Carba1 and sub-therapeutic doses of PTX elicits anti-tumor activity in xenograft tumor-bearing mice (*12*). We posited that this combination may also mitigate the adverse effects associated with higher doses of PTX, such as those promoting CIPN. To investigate this, we directly tested the effect of Carba1 on models of PTX-associated CIPN. In patients with CIPN, a hallmark symptom is altered mechanical sensitivity resulting from the preferential damage to primary afferent long sensory nerve fibers, with or without accompanying demyelination (*20*, *21*). Dying-back axonopathy is observed in both patients with CIPN and rodent models, establishing distal-to-proximal axonal peripheral nerve degeneration as a key factor underlying CIPN development (*10*, *22*). First, we examined whether Carba1 could prevent the axonal degeneration of sensory neurons exposed to pathogenic doses of PTX (Fig. 2). To this end, primary cultures of sensory neurons from dissociated adult mouse dorsal root ganglia (DRGs) were grown in culture for 12 days, allowing them to fully recover from axotomy and extend their axons. The cultures were then exposed to Carba1 at a dose (12 µM) which alone did not show significative effects on neurite fragmentation, with or without PTX (50 nM) administered for 72 h. This concentration and timing of PTX treatment are widely reported to induce axonal degeneration that phenocopies the *in vivo* pathology (*23*, *24*). Neurons were then fixed and stained for neurofilament to assess the extent of axonal fragmentation by calculating the proportion of the total axonal area covered by the fragmented axons. As expected, we found that PTX induced robust axonal degeneration under these experimental conditions (Fig. 2A, B). However, co-treatment of neurons with Carba1 and PTX prevented neuronal degeneration, as shown by the reduced number of fragmented axons (Fig. 2A, B), indicating that Carba1 offers neuroprotection against PTX-induced axonal degeneration.

**Fig. 2.**
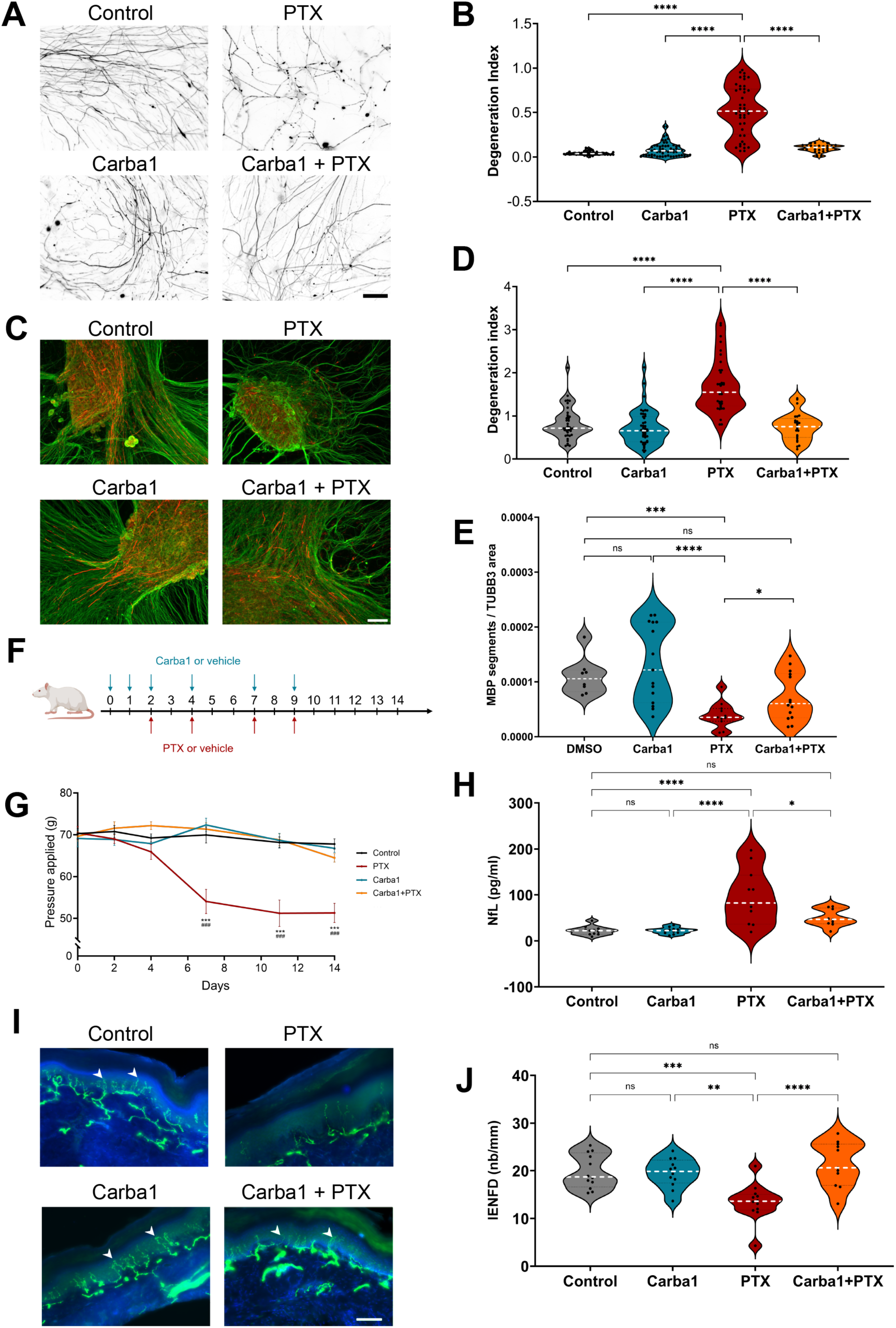
Carba1 prevents PTX-induced neuropathy *in vitro* and *in vivo*. **(A)** Representative images of neurofilament staining in axons of adult mouse DRG neurons at 12 days *in vitro* (DIV) treated for 72 hours with Dimethylsulfoxide (DMSO, Control), 50 nM of paclitaxel (PTX), 12 µM of Carba1or their combination as indicated. Scale bar, 50 μm. **(B)** Quantification of the effect of the different treatments on axonopathy, by the degeneration index. Data are pooled from 3 experiments (n = 16 to 55 neurites per condition for each experiment). **** p<0.0001, ANOVA. **(C)** Representative images of mouse dorsal root ganglia (DRG) explants (DIV 14) at 7 days in differentiation medium to allow Schwann cell myelination. DRG explants were treated for 72 hours with DMSO (Control), 12 µM of Carba1, 500 nM of paclitaxel (PTX), or their combination as indicated. Neurons were stained with anti-β3-tubulin (TUBB3, green) and anti-myelin basic protein (MBP, red) antibodies. Scale bar, 100 µm. **(D)** Quantification of the effect of the different treatments on axons of DRG explants, as assessed by the degeneration index calculated as explained in the material and methods section. **** p<0.0001, ANOVA. **(E)** Quantification of the effect of the different treatments on myelination, represented as the ratio between the sum of the number of MBP segments and the axonal (TUBB3) area, calculated from 5 random fields per DRG explant from at least 3 different experiments. * p = 0.018, *** p<0.001, Mann-Whitney test. **(F)** Illustration of the experimental design for the rat model of PTX-induced neuropathy. Carba1 (50 mg/kg, blue arrows) was injected two days (day 0 and 1) before PTX (5 mg/kg, red arrows) administrations in combination or not with Carba1 at the indicated time points. **(G)** Effects of Carba1 on tactile allodynia in a rat model of PTX-induced neuropathy. Behavioral assessments were performed at day 0 (basal values) and at day 2, 4, 7, 11 and 14. Control animals (black curves) received vehicle injections. The response of the animals injected with Carba1 (blue curve), PTX (red curve) and Carba1 together with PTX (orange curve) are shown at the indicated times. The results are presented by the mean ± SEM (n = 12 per group); *** p < 0.001 PTX *vs.* Control; ### p < 0.001 PTX *vs* Carba1+PTX, ANOVA for repeated measures followed by a *post hoc* Tukey test. **(H)** Concentrations of neurofilament light chain (NfL) in rat blood samples collected on day 15, before euthanasia of rats used in the experiment shown in F/G. * p=0.014, **** p<0.0001, ANOVA. **(I)** Representative confocal images of intraepidermal nerve fibers (IENFs) from fixed rat hindpaw biopsies collected on day 15. IENFs (white arrows) were immunolabeled with PGP9.5 (green) and project from subepidermal fascicles across the epidermal-dermal junctions. Scale bar, 100 µm. **(J)** Quantification of IENF densities from hind paw biopsies. Diagrams represent mean ± SEM calculated from individual rat IENF densities from each group (black points), n = 12 rats/group. ** p<0.01, *** p<0.001, ****p<0.0001, ns non-significant, ANOVA.

DRG explant cultures have the advantage of preserving the original architecture of the DRG, maintaining the relationships between neurons, Schwann cells (SC), and fibroblasts. We thus examined the effect of Carba1 on PTX-induced SC injury by using *ex vivo* cultures of mouse DRG explants, which initiate to myelinate their axons starting from two weeks in culture. We observed that 500 nM PTX was required to induce neuronal degeneration in DRG explants, most likely a result of the multicellular nature of these 3D structures, which can influence the diffusion and absorption of the drug. We noticed that under these conditions, PTX also affected the overall structure of DRGs, which appeared smaller and with a reduced axonal network density, compared to controls (Fig. 2C, D). However, when DRG explants were treated with both PTX and Carba1 (12 µM) the global structure of the DRGs and the neuronal network density were like the control group (Fig. 2C), confirming the neuroprotective effect of Carba1 also in the explants (Fig. 2D, fig. S1A).

We performed Myelin Basic Protein (MBP) staining to assess axonal myelination and found that while Carba1 alone had no effect, PTX caused a significant reduction in MBP staining. However, when Carba1 was combined with PTX, myelin staining was substantially improved compared to PTX alone (Fig. 2C, fig. S1B) and quantification confirmed that this increase was statistically significant (Fig. 2E).

Next, we set out to evaluate the neuroprotective efficacy of Carba1 in a rat model of CIPN. Carba1 (50 mg/kg) was injected intraperitoneally 2 days (day 0 and day 1) prior to the initiation of PTX treatment (5 mg/kg) and repeated with each PTX injection. Rats then received either PTX, Carba1 or a combination of both on day 2, 4, 7 and 9 (Fig. 2F). The cumulative dose of PTX was 20 mg/kg, that of Carba1 was 300mg/kg, administrated either alone or in combination. A behavioral test to assess mechanical allodynia (electronic Von Frey test), characteristic of CIPN (*25*), was carried out on days 0, 2, 4, 7, 9, 11 and 14. No significant difference in the weight of the animals was observed between the different groups (fig. S1C), indicating that neither treatment had a detectable negative impact on the general health of the animals. However, rats treated with PTX developed a tactile allodynia with a significant decrease of paw withdrawal thresholds at day 7 (p < 0.001), in comparison to basal value at day 0, and this allodynia was still present 5 days after the last injection of PTX (Fig. 2G). There was no difference between the tactile threshold of animals treated with Carba1 alone and that of the control group, indicating that Carba1 *per se* was not analgesic. However, rats treated with the combination of Carba1 and PTX did not differ from the controls and their response threshold was significantly different from that of PTX (p < 0.0001) (Fig. 2G), indicating that Carba1 prevents PTX-induced tactile allodynia.

We then investigated whether this protective effect of Carba1 was also detectable by histopathological analysis of intraepidermal nerve fiber density (IENFD) and serum concentration of neurofilament light chain (NfL), a biomarker of PTX-induced peripheral neuropathy (*25*, *26*). To that end, skin biopsies of hind paws collected from sacrificed animals were examined for IENFD assessment and blood samples collected for NfL analysis.

We found that the NfL serum concentration was significantly increased by PTX treatment compared to control and Carba1 treatment. When Carba1 and PTX treatments are combined, however, the NfL serum concentration was significantly reduced compared to PTX treatment (Fig. 2H).

We observed that PTX also significantly reduced IENFD by 30% whereas Carba1 had no effect and was not different from controls. However, co-treatment with Carba1 prevented PTX-induced IENF degeneration confirming its neuroprotective effect at the histopathological level (Fig. 2I, J).

### Carba1 also prevents cisplatin- and bortezomib-induced neurotoxicity in adult DRG neurons

In addition to MT-targeting anticancer drugs like PTX, several other classes of anticancer agents with distinct antineoplastic mechanisms of action can induce CIPN. We thus explored if Carba1 could also protect against neuronal degeneration promoted by either the DNA-binding drug Cis or the proteasome inhibitor and MT-stabilizing drug Bort.

To this end, we compared the effect of these two anticancer drugs, with or without Carba1 on primary cultures of neurons from dissociated adult mouse DRG as we did for PTX (Fig. 2A, B). Exposing the neurons for 72 h to 10 µM Cis or 100 nM Bort induced significant axonal fragmentation assessed by staining the axons with the neurofilament antibody (Fig. 3A).

**Fig. 3.**
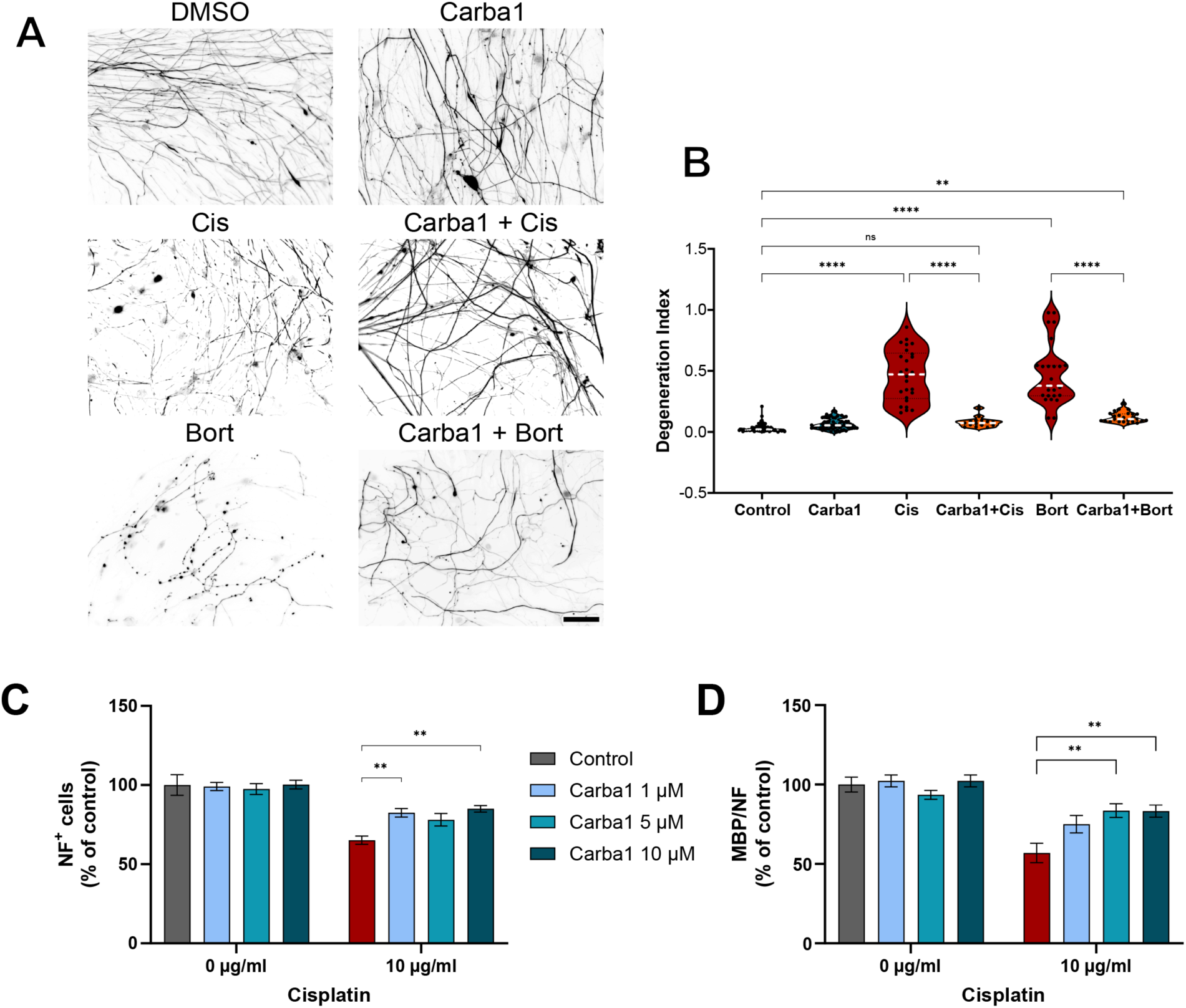
Preventive effect of Carba1 on cisplatin- and bortezomib-induced neuropathies. **(A)** Representative images of neurofilament staining in axons of adult mouse DRG neurons (DIV 12) treated for 72 h with DMSO (Control), 12 µM of Carba1, 10 µM of Cisplatin (Cis), 100 nM of Bortezomib (Bort), or a combination of the drugs with Carba1 as indicated. Scale bar, 50 μm. **(B)** Quantification of the degree of axonopathy in DRG neurons treated as in (**A**) by degeneration index. Data are pooled from 3 independent experiments (8 fields per condition for each experiment). ** p<0.01, **** p<0.0001, ANOVA. **(C)** Quantification of the effect of DMSO (Control) or Cisplatin (Cis, 10 µg/ml) in combination with 1 µM, 5 µM, 10 µM Carba1 on the number of rat embryonic DRG neurons. ** p<0.01, ANOVA. **(D)** Quantification of the degree of myelination in rat embryonic DRG neurons treated as in (**C**) and stained for MBP. ** p<0.005, ANOVA.

However, co-treatment with Carba1 (12 µM) was able to prevent neuronal fragmentation induced by either Cis or Bort (Fig. 3A). Axonal degeneration was quantified using the degeneration index and showed that Cis and Bort both significantly increased the degeneration index. When Cis or Bort were co-administrated with Carba1, the value of the index was close to the control value (Fig. 3B).

To evaluate the Carba1protective effect on neuronal myelination, we analyzed the impact of increasing doses of Carba1 on dissociated embryonic DRG neurons exposed to a toxic dose of Cis (10 µg/mL). We found that Carba1 exhibited its maximal protective effect against Cis-induced demyelination at 5 µM (Fig. 3C, D). These results indicate that Carba1 is also able to prevent Cis- and Bort-induced neurotoxicity, by protecting axonal integrity and myelination.

### Carba1 regulates energy metabolism through direct binding and activation of NAMPT

Given that Carba1 protects neurons from degeneration induced not only by PTX but also by drugs with different mechanisms of action, we hypothesized that its neuroprotective activity was due to activation of a target other than the taxanes site of tubulin, with the underlying mechanism being generally beneficial to neuronal health.

Neurons are among the most energy-demanding cell types and bioenergetic failure is considered one of the main contributors to neuronal degeneration (for review see (*27*)). Therefore, we first conducted an unbiased comparative metabolomic profiling of Carba1 treatment using proton-based solution nuclear magnetic resonance (1H NMR). Given the generality of the metabolic pathways and to gain sensitivity and reproducibility, we carried out this experiment on HeLa cells rather than isolated neurons. Aqueous extracts of HeLa cells, treated or not with 12 µM Carba1 for 2 hours were analyzed by NMR. Multivariate analyses, Orthogonal Projections to Latent Structures-Discriminant Analysis (OPLS-DA), significantly separated the metabolic fingerprints of each condition into two distinct groups (Fig. 4A, fig. S2). The metabolites responsible for the changes in the metabolic profile of Carba1-treated samples compared to the control are illustrated on the loading plot (Fig. 4B). Overall, this metabolic signature shows an enhanced energetic metabolism involving glycolysis (increased lactate), glutaminolysis (increased glutamate and low glutamine), and increased ATP, creatine and phosphocreatine levels. Univariate statistical analysis to quantify the mean relative amplitude for each metabolite revealed a significant increase in GTP levels. Interestingly, such univariate analysis also uncovered a significant decrease of NAD^+^ upon Carba1 treatment (Fig. 4C). These results indicate that Carba1 modulates energy metabolism. However, NADH levels were below the limit of reliable quantification, preventing further estimation of the NAD^+^/NADH balance.

**Fig. 4.**
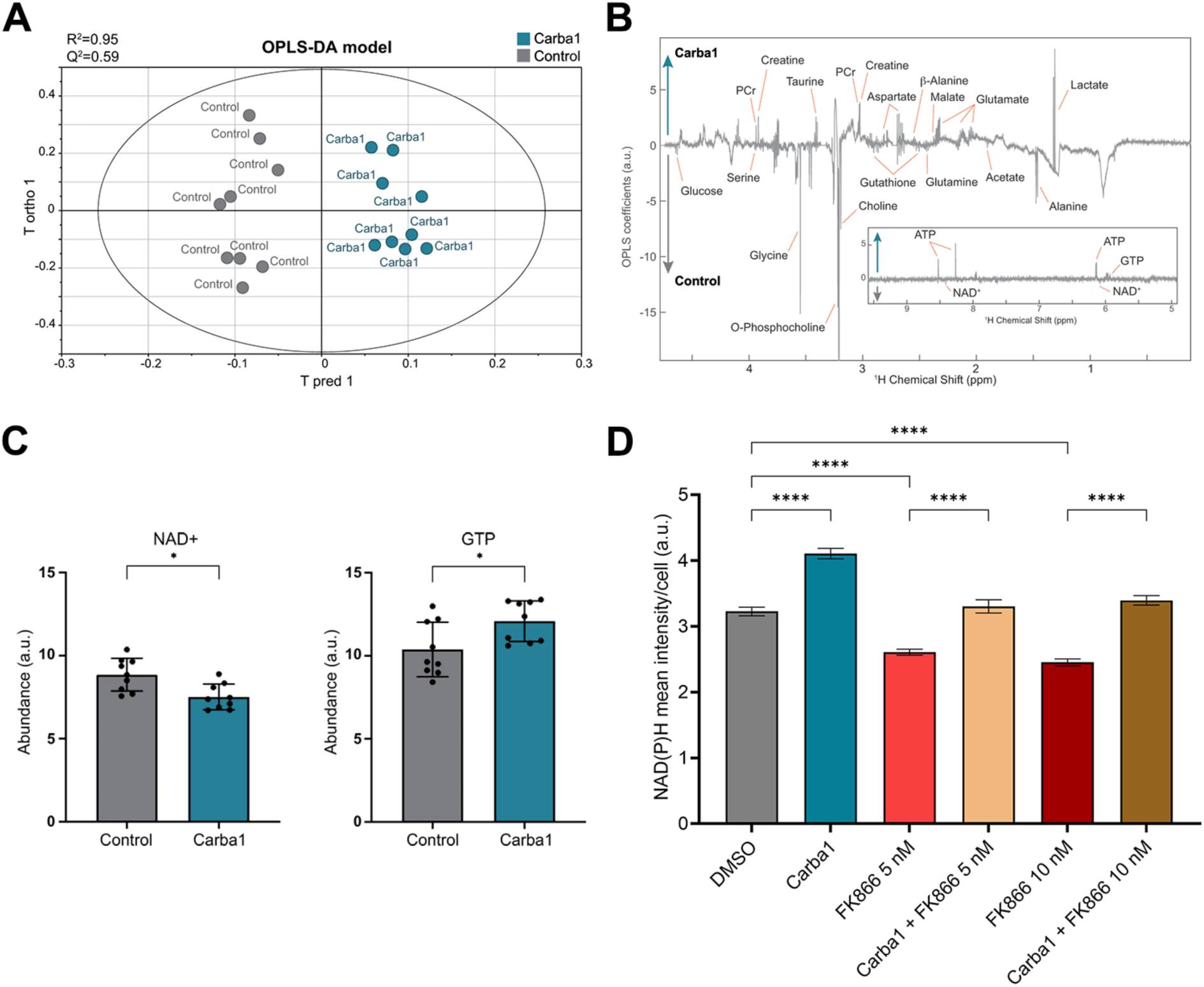
Effect of Carba1 on cell metabolism. **(A)** Score plot from OPLS-DA multivariate modelling of the NMR metabolic profiles of cells treated for 2 h with 12 µM Carba1 (blue points) or DMSO (Control, grey points). N=18, with 9 replicates per sample class; 1 predictive +3 orthogonal components; R^2^Y=0.95; Q^2^=0.59). **(B)** Associated OPLS-DA back-scaled loadings. Insert: same plot, additional region [5, 9.5 ppm] **(C)** Effect of Carba1 treatment on the abundance of NAD^+^ (left) and GTP (right). * p< 0.05, t test. **(D)** Measurement of NAD(P)H accumulation in HeLa cells. The intensity of the NAD(P)H autofluorescent signal was quantified in living cells and normalized to the cell area. Data represent the mean ± SEM of 3 independent experiments, with a minimum of 40 cells per condition.**** p<0.0001, ANOVA.

To find out whether Carba1 also affected cellular NADH production, we measured NAD(P)H autofluorescence following ultraviolet excitation, a minimally invasive optical approach (*28*). Live HeLa cells were incubated for 24 h with 12 µM Carba1 and NAD(P)H intensity was monitored by confocal microscopy. We observed a significant increase of about 30% in NAD(P)H production by cells treated with Carba1 (Fig. 4D).

We posited that the metabolic changes observed (decrease in NAD^+^, increase in NADH) could be explained by the influence of Carba1 on several molecular targets involved in the regulation of NAD^+^/NADH levels and energy metabolic pathways such as, for example, inhibition of sterile alpha and TIR motif-containing 1 enzyme (SARM1) (*29*, *30*), inhibition of poly(ADP-ribose) polymerase (PARP) (*31*) or activation of nicotinamide phosphoribosyltransferase (NAMPT) (*32*). NAMPT is the rate-limiting enzyme in the NAD^+^ salvage pathway that converts nicotinamide (NAM) to nicotinamide mononucleotide (NMN), which is responsible for most of the NAD^+^ formation. We initially investigated whether NAMPT might be a target of Carba1. This was prompted by Carba1’s carbazole core, which is similar to P7C3 (fig. S3) - a compound believed to target NAMPT (*33*) - as well as by evidence that NAMPT activators have demonstrated neuroprotective effects (*34*, *35*). To this end, we used the potent NAMPT inhibitor FK866 (*36*), which is considered the pharmacological equivalent of genetic NAMPT silencing for NAMPT target validation (*37*).

We observed that a 24-hour incubation of cells with 5 and 10 nM FK866 was indeed able to reduce NAD(P)H production by approximately 20%. When cells were pre-incubated with 12 µM Carba1 and then co-treated with 5 or 10 nM of FK866, the levels of NAD(P)H were similar to the control levels, indicating that Carba1 was able to counteract the FK866 effect on NAD(P)H production, at both concentrations of FK866 (Fig. 4D).

FK866-induced cell depletion of NAD^+^/NAD(P)H can lead to cell death over several days. We thus investigated if Carba1 could relieve the cytotoxicity mediated by FK866 in HeLa cells, and compared Carba1 with P7C3 and the recently described potent activator of NAMPT, NAT (*34*). As shown in Fig. 5A, FK866 induced HeLa cell death, with a GI_50_ of 2.4 ± 0.09 nM. However, co-administration of Carba1 or NAT with FK866 counteracted the toxic effect of FK866 in a dose-dependent manner, inducing a shift to the right of the viability curves (GI_50_: 6.5 ± 1.3 nM for 12 µM Carba1, and 7.2 ± 2.4 nM for 12 µM NAT). Interestingly, P7C3 was unable to counteract the toxic effect of FK866. In contrast, Carba1’s ability to counteract FK866-induced cell death strongly suggests that, like NAT, Carba1 binds to NAMPT.

**Fig. 5.**
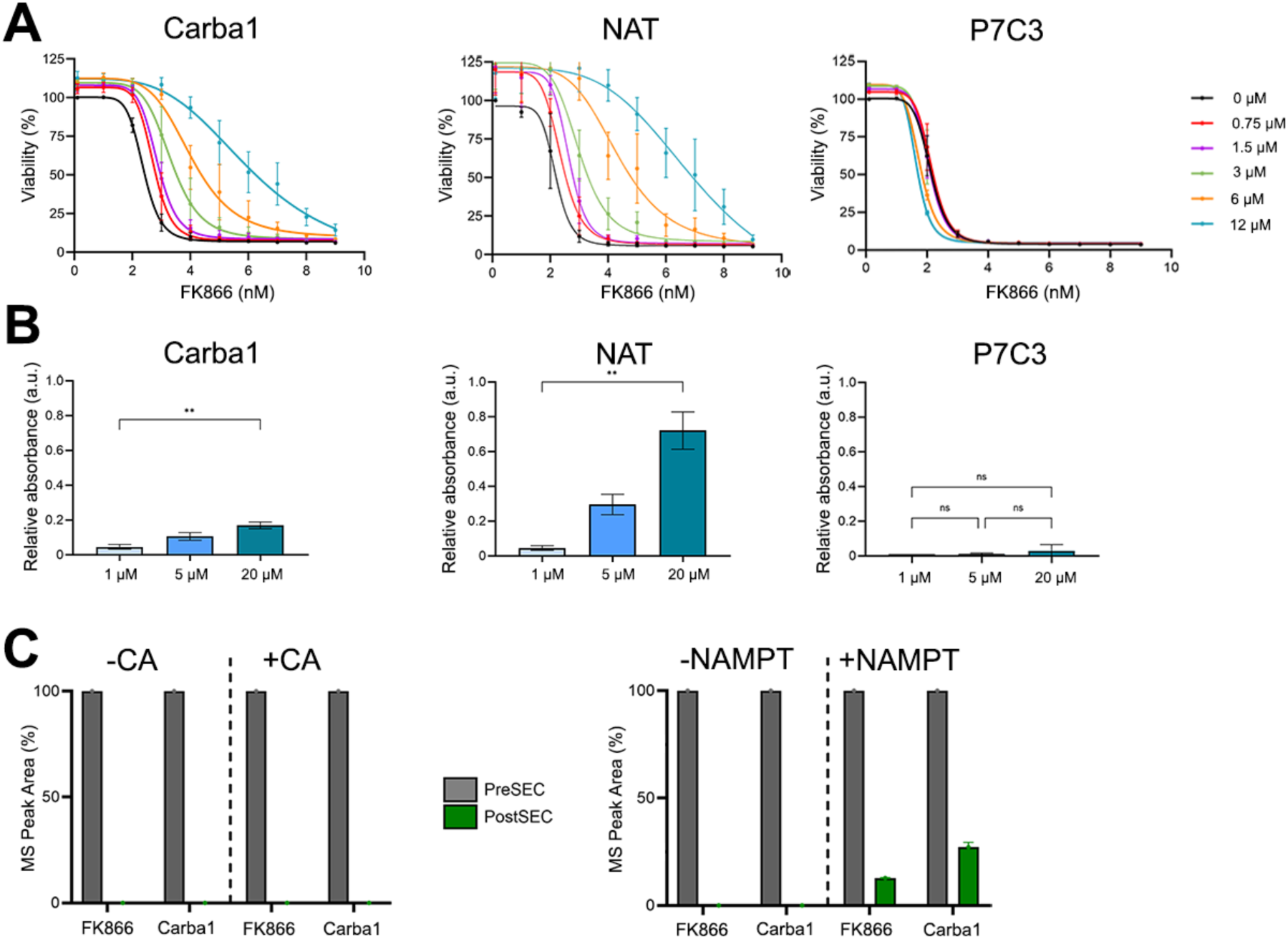
Carba1 targets NAMPT. **(A)** Carba1 and NAT, but not P7C3 relieve the cytotoxicity caused by the NAMPT inhibitor FK866 on HeLa cells. Viability dose-effect curves of FK866 in the presence of increasing doses of Carba1 (left), NAT (middle) and P7C3 (right). Data represent the mean ± SEM from 3 independent experiments. **(B)** *In vitro* dose-dependent activation of NAMPT by Carba1 and NAT but not P7C3. A triply coupled NAMPT assay was performed in the presence of the indicated doses of the compounds. Data obtained after 60 min of incubation were normalized by subtraction of their respective DMSO value, which corresponds to the basal activity of NAMPT. ** p<0.005, ANOVA. **(C)** NAMPT binding assay using AS-MS. 10 µM Carba1, were incubated with 3 µM NAMPT or 3 µM Carbonic Anhydrase (CA, negative control). 10 µM FK866 was assayed as positive control of NAMPT binding. Protein–ligand complexes were then separated from the unbound compounds by size-exclusion chromatography (SEC) and the amount of protein-bound compounds quantified by mass spectrometry after dissociation from the protein. Histograms show the amounts of compound before (preSEC, grey) and after (postSEC, green) size exclusion chromatography (SEC) in the presence of CA or NAMPT.

We next tested whether Carba1 could activate NAMPT *in vitro*. To that aim, we used a commercial recombinant human enzyme assay containing four enzymes: NAMPT, nicotinamide nucleotide adenylyltransferase 1 (NMNAT1), alcohol dehydrogenase (ADH) and diaphorase (see methods section). In the presence of NAM and 5-phosphoribosyl-1-pyrophosphate (PRPP), NAMPT converts NAM to NMN, which is then converted into NAD^+^ by NMNAT1, while ADH converts NAD^+^ to NADH. Then, into a coupled-reaction, diaphorase converts NADH into NAD^+^ and tetrazolium salt WST-1 (4-[3-(4-iodophenyl)-2-(4-nitro-phenyl)-2H-5-tetrazolio]-1,3-benzene sulfonate) to formazan, which can be detected at OD_450_ nm. In practice, we proceeded in two steps. First, Carba1 and the different tested compounds were incubated for 15 minutes with NAMPT, NMNAT1, NAM and PRPP to allow NAD^+^ production. The reaction was then stopped with FK866 before adding ADH and diaphorase, to convert the NAD^+^ produced to NADH, and absorbance was monitored at regular intervals. Fig. 5B shows the 60-minute values for Carba1, NAT and P7C3, normalized by subtraction of their respective DMSO value, which corresponds to the basal NAMPT activity. We found that Carba1 enhanced NAMPT activity in a dose-dependent manner, similar to NA, although less potent. No significant effect of P7C3 was observed in this assay (Fig. 5B).

We then tested whether Carba1 directly binds to NAMPT, using affinity selection-mass spectrometry (AS-MS) (*38*, *39*). To this end, Carba1 or FK866 (positive control of NAMPT binding) were first incubated with NAMPT or carbonic anhydrase (CA, negative control). Protein– ligand complexes were separated from the unbound compounds by size-exclusion chromatography (SEC). To quantify the protein-bound fraction, the complexes were then dissociated using reverse-phase chromatography and the compounds were analyzed and quantified by mass spectrometry. By comparing the amounts of compound before (preSEC) and after (postSEC) size exclusion chromatography in the presence of CA or NAMPT, we assessed their ability to selectively bind to NAMPT. As shown on Fig. 5C, neither FK866 nor Carba1 were found to bind CA. However, both compounds were recovered in the fraction containing NAMPT, indicating that like FK866, Carba1 directly binds to NAMPT.

### Structure-activity relationship (SAR) analysis of the carbazole series enables the identification of moieties essential for the synergistic activity and those involved in the metabolic activity

Our studies indicate that Carba1 has two primary targets: it not only targets tubulin and synergizes with taxanes, but it also activates the NAMPT enzyme, thus providing neuroprotective effects in neurons affected by chemotherapeutic agents.

To gain insights into how Carba1 (Fig. 6A) interacts with these targets and whether the same chemical moieties are involved in these interactions, we conducted a structure-activity relationship (SAR) analysis on 16 structural analogs of Carba1. The positions of Carba1 targeted for SAR investigation are shown in Fig. 6B.

**Fig. 6.**
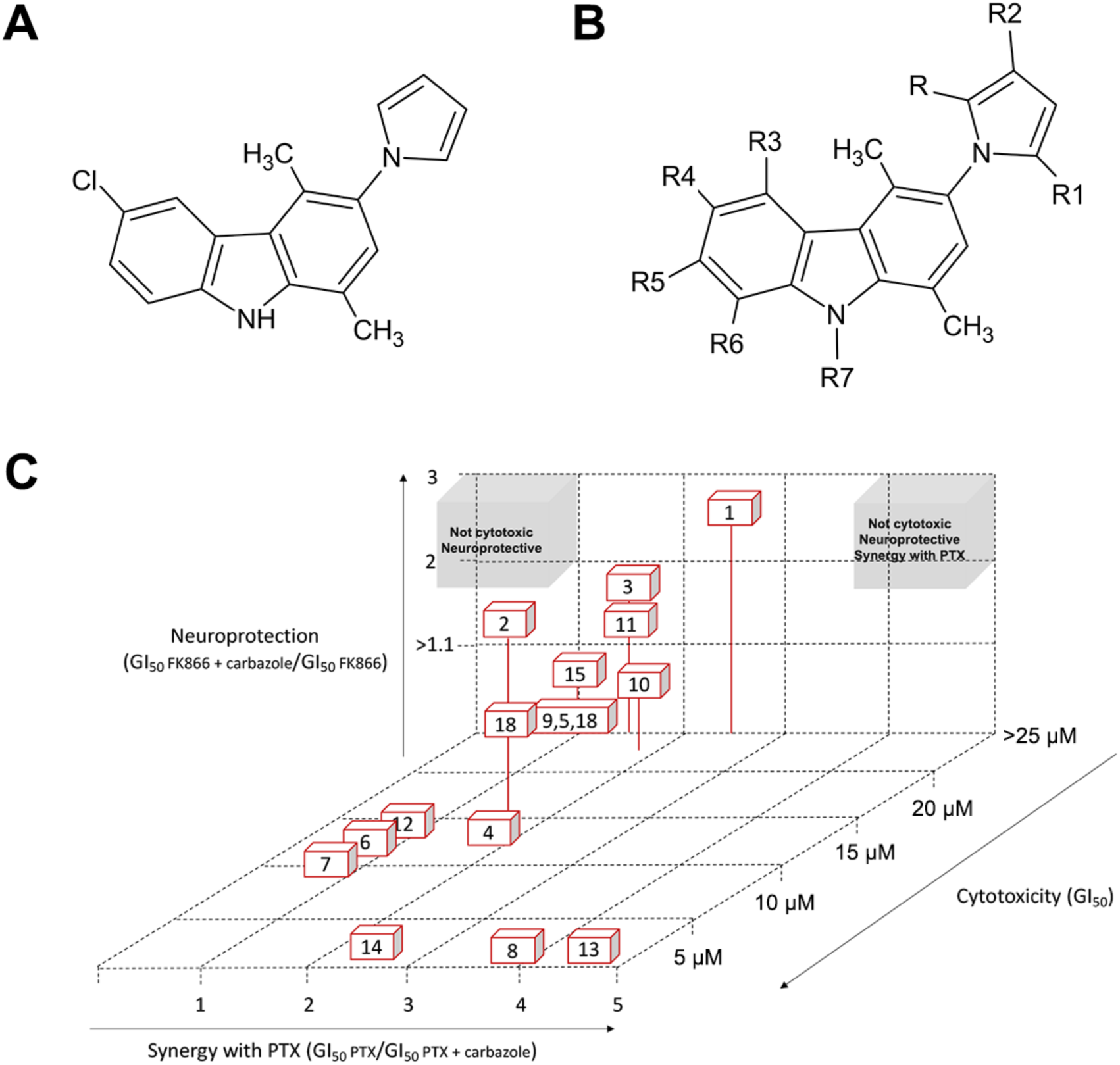
Structure activity relationship analysis of Carba1. **(A)** Carba1 chemical structure **(B)** Positions targeted for SAR investigation **(C)** Compared activities of Carba1 (1) and its analogs.

We conducted the SAR analysis by comparing the different cellular activities of these compounds. We first analyzed the effect of each compound on cell viability. We found that substituting R2, with a hydroxymethyl or a formyl group, conferred higher cytotoxicity (GI_50_ in the micromolar range) to compounds **8** and **13** respectively, compared to Carba1 (**1**). Similarly, the substitution of R2 by a formyl group of the bromo analog of Carba1, compound **2**, induced a higher cytotoxicity to compound **14** (Table 1 and Fig. 6B). The substitution of R2 by a methylamino group also made compounds **6** and **7** more toxic than Carba1, but to a lesser extent (GI_50_ in the decamicromolar range). The same applies to the positional isomer of Carba1, compound **4** with a chlorine atom in R5, and also to the dimethylpyrrolyl analog of Carba1 (**12**). The dimethyl analog **9** of the bromo derivative **2**, however, was devoid of the cytotoxicity expressed by the latter (Table 1).

**Table 1:** SAR of Carba1 substitution. * The ratios were calculated for the highest non cytotoxic dose of carbazole analogs

| Compound | R | R1 | R2 | R3 | R4 | R5 | R6 | R7 | GI<sub>50</sub> (μM) | GI<sub>50</sub> PTX /<br>GI<sub>50</sub> PTX with carbazole * | GI<sub>50</sub> FK866 with carbazole /<br>GI<sub>50</sub> FK866 * |
| --- | --- | --- | --- | --- | --- | --- | --- | --- | --- | --- | --- |
| 1 | H | H | H | H | Cl | H | H | H | > 25 | 2.5 | 2.6 |
| 2 | H | H | H | H | Br | H | H | H | 16 | 1.6 | 2 |
| 3 | H | H | H | H | H | H | Et | H | > 25 | 1.5 | 1.5 |
| 4 | H | H | H | H | H | Cl | H | H | 15 | 1.8 | 0.93 |
| 5 | H | H | H | Cl | H | H | H | H | > 25 | 1 | 1.07 |
| 6 | H | H | CH<sub>2</sub>NMe<sub>2</sub> | H | Cl | H | H | H | 12.7 | 0.8 | 1 |
| 7 | H | H | CH<sub>2</sub>NHEt | H | Cl | H | H | H | 10.3 | 0.77 | 1 |
| 8 | H | H | CH<sub>2</sub>OH | H | Cl | H | H | H | 1 | 3.8 | 1 |
| 9 | Me | Me | H | H | Br | H | H | H | > 25 | 1 | 0.99 |
| 10 | H | H | H | H | OC(O)OEt | H | H | H | 25 | 1.7 | 1.1 |
| 11 | H | H | H | H | OH | H | H | H | > 25 | 1.5 | 1.3 |
| 12 | Me | Me | H | H | Cl | H | H | H | 14 | 0.9 | 1.09 |
| 13 | H | H | CHO | H | Cl | H | H | H | 1 | 4.4 | 1 |
| 14 | H | H | CHO | H | Br | H | H | H | 1.2 | 2.5 | 1 |
| 15 | Me | Me | H | H | OC(O)OEt | H | H | H | > 25 | 1 | 1.1 |
| 16 | H | H | H | H | Cl | H | H | CH<sub>2</sub>CH(F)CH<sub>2</sub>NH-3-MeOPhe | > 25 | 1.2 | 0.98 |

Using the highest non-cytotoxic dose of each compound, we quantified the potential for synergy with PTX by the ratio GI_50_ PTX / GI_50_ (compound + PTX), and the potential for activation of NAMPT by the ratio GI_50_ (compound + FK866)/ GI_50_ FK866 (Table 1).

For some compounds, there is a strong correlation between the cytotoxicity of the compounds and their ability to act synergistically with PTX. The more cytotoxic compounds, **13**, **8** and **14** showed also the best synergistic activity with PTX. The relatively less cytotoxic compounds **2** and **4** are also less likely to act synergistically with PTX. Conversely, the moderately cytotoxic derivatives **6**, **7** and **12** are devoid of a synergistic potential with PTX. The other most noticeable exception is Carba1, which is poorly cytotoxic (GI_50_ higher than 25 µM) while showing a good synergistic activity, allowing reduction of PTX cytotoxic doses by a factor of 2.5. Compound **10**, which carries an ethylcarbonate group instead of the chlorine atom of Carba1, is also poorly cytotoxic, but still shows a good synergistic activity, although less noteworthy than that of Carba1.

Using the resistance to FK866 cytotoxic activity as a read-out, only four compounds, including Carba1, were found able to activate NAMPT (ratios GI_50_ FK866 with carbazole/ GI_50_ FK866 >1.1). The presence of a halogen at R4, associated with an H at R and R1, appears to be important for the NAMPT activating activity, since the bromo analog **2** of Carba1 is still quite neuroprotective even if it becomes more cytotoxic. A low level of NAMPT activating activity is nevertheless retained if the halogen in R4 is replaced by a hydroxyl group (**11**) or if the halogen is removed, while an ethyl group is carried in R6 (**3**). The ethylcarbonate compound **10** is still very slightly neuroprotective, perhaps due to its hydrolysis into the hydroxyl derivative **11**. The dimethyl analog of **10**, compound **15**, is also poorly neuroprotective. All types of activity are lost when the chlorine atom of Carba1 moves from R4 to the R3 position (**5**) or when the pyrrolyl ring of **2** is further dimethylated (**9**).

As Carba1 shares a carbazole scaffold with P7C3, but differs mainly by the addition of a - CH2CH(F)CH2NH-3-MeOPhe on the central ring nitrogen (fig. S3), we substituted the hydrogen on the central ring nitrogen of Carba1 with -CH2CH(F)CH2NH-3-MeOPhe (**16**). We found that such a substitution resulted in the loss of all types of activity, underlining the crucial role of this moiety.

Finally, no correlation was found between the cytotoxic/synergistic activities of the compounds and their ability to activate the NAMPT enzyme, since the more cytotoxic compounds, which are also the compounds that synergize the best with PTX (**13**, **8** and **14**), are devoid of neuroprotective activity. This SAR study indicates that it is possible to develop compounds with either synergistic or neuroprotective activity, or both, while being devoid of cytotoxicity. Pharmacomodulations of Carba1 (**1**) will be pursued with the objective to obtain derivatives located in the upper right part of Fig. 6C, or only neuroprotective compounds, located in the upper left part of Fig. 6C.

### Carba1 does not interfere with tumor growth or PTX anti-cancer activity

As we have identified that Carba1 prevents CIPN through NAMPT activation, an important consideration is whether Carba1 could accidentally promote tumor growth or interfere with therapeutic doses of PTX in mice. Our previous *in vivo* experiments, designed to analyze the synergistic activity of Carba1 with low doses of PTX, ruled out the possibility that Carba1 could have a pro-tumoral activity (*12*). Nonetheless, we herein designed an *in vivo* experiment to specifically test the effect of a high dose of Carba1 on tumor growth and anti-cancer activity when applied alone or in combination with an antitumor dose of PTX that overrides the synergistic effect of Carba1 on tumor growth inhibition. Immunodeficient nude mice were injected subcutaneously in the flank with exponentially dividing HeLa cells to induce tumors. Once the tumors were established (> 120 mm3), mice received intravenous injections of either PTX (8 mg/kg), Carba1 (60 mg/kg), a combination of PTX (8 mg/kg) and Carba1 (60 mg/kg) or vehicle alone every 2 days for a duration of 10 days. All mice showed a slight body weight loss after the first injection regardless of the treatment, but then their weight increased again and reverted to normal except for one out of the eight mice of the vehicle group (fig. S4).

All clinical data analyzed in this study (blood counts, markers of renal or hepatic function) indicated that none of the treatments had any major toxicological effects (fig. S5). More importantly, while we found that PTX administration drastically reduced tumor size, there were no significant tumor size changes by co-treatment with Carba1 in either vehicle-treated mice or PTX-treated mice (Fig. 7A, B). This suggests that Carba1 neither promotes tumor growth nor inhibits the antitumor activity of an effective dose of PTX.

**Fig. 7.**
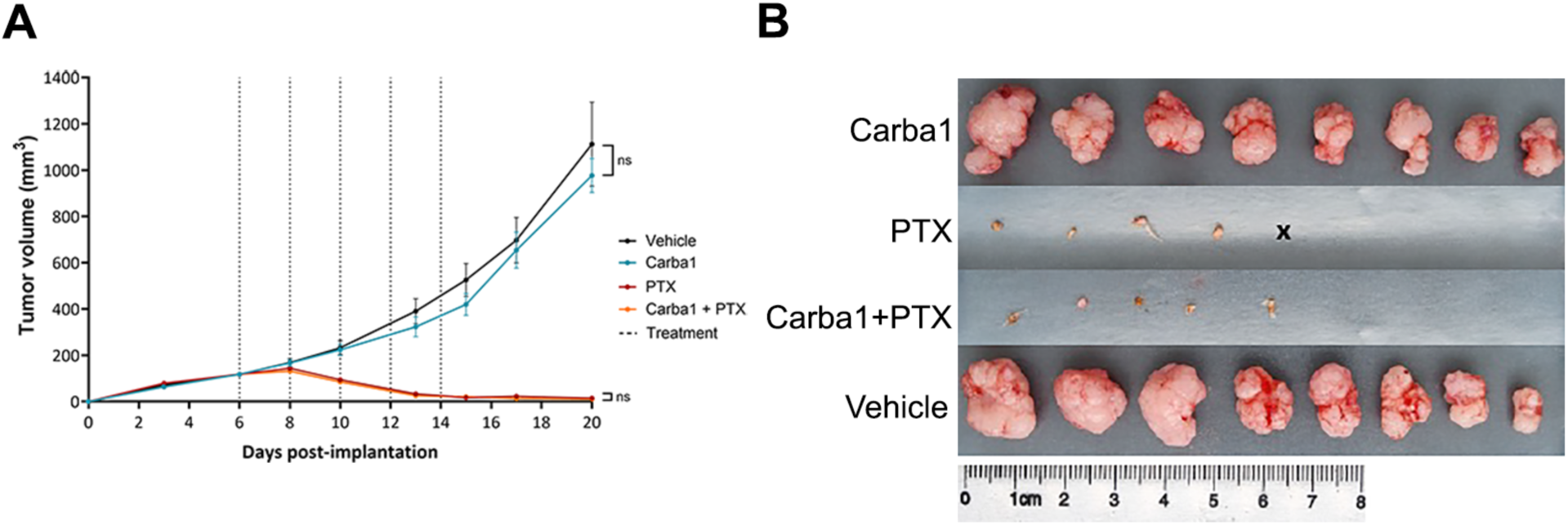
Carba1 does not affect antitumoral efficacy of a therapeutic dose of PTX. **(A)** Analysis of the effect of paclitaxel (PTX), Carba1, and their combination on the growth of HeLa cells xenografted in mice. HeLa cells were subcutaneously implanted on the flank of female athymic nude mice. When the tumors reached a volume of about 120 mm^3^, mice were treated every two days (dotted lines) with PTX (8 mg/kg), Carba1 (60 mg/kg), the combination of PTX (8 mg/kg) and Carba1 (60 mg/kg), or vehicle. Tumor growth was monitored with a sliding caliper. Data are mean ± SEM, n = 8 mice per group, ANOVA. **(B)** Images of the tumors isolated at the end of the experiment.

## DISCUSSION

Advances in cancer treatments have improved patient survival but revealed persistent side effects like chemotherapy-induced peripheral neuropathy (CIPN). Unlike other adverse effects, CIPN can endure long term, often requiring dose reductions or treatment discontinuation, which jeopardizes clinical outcomes. While symptomatic pain relief exists, no effective strategies currently prevent CIPN.

We describe Carba1, a novel bi-functional agent that protects peripheral neurons from chemotherapy-induced toxicity while lowering required doses of taxanes. *In vitro*, Carba1 synergizes with compounds targeting the tubulin taxane site, disrupting microtubule dynamics to enhance the action of sub-saturating doses of PTX. Additionally, Carba1 combats the neurotoxicity of CIPN-inducing agents by activating NAMPT, a key enzyme in the NAD^+^ salvage pathway, supporting a common neuroprotective mechanism.

We demonstrate *in vitro* that Carba1 synergizes with compounds targeting the tubulin taxane site, but not with other chemotherapeutic agents, Cis or Bort, with different mechanisms of action. This further supports the conclusion that Carba1 exerts its effect by perturbing MT dynamics at the growing end, promoting the binding of non-saturating doses of PTX (*12*, *40*).

Although PTX, Cis or Bort have different anti-tumor mechanisms of action, their main non-hematological adverse effect is CIPN. The mechanisms responsible for this “dying-back” of distal nerve endings are not completely understood and it is unknown whether distinct agents engage in different cell death programs or converge on a shared axon degeneration pathway (*41*–*43*). However, at least two drugs with distinct mechanisms of antitumor action, vincristine and Bort, were reported to implement distinct upstream mechanisms to activate a common sterile alpha and TIR motif containing 1 (SARM1)-dependent axon degeneration program that converge on axonal NAD^+^ depletion (*44*). Here, we demonstrate that Carba1 protects against the neurotoxicity of three different classes of CIPN-compounds through the binding and activation of the enzyme NAMPT, the rate-limiting enzyme in the NAD^+^ salvage pathway, supporting a common mechanism of neuroprotection.

An impairment of myelinated fiber function could be responsible for the sensory disturbances in CIPN (*45*–*47*). Carba1 also exerts a protective effect on Schwann cells, which produce the myelin sheath, significantly sparing them from the toxicity of PTX or Cis. These results support a general mechanism of neuroprotective action of Carba1, which might be effective not only in toxicity-induced, but also in related types of familial neuropathies.

Carba1 demonstrates neuroprotective efficacy in a rat model of PTX-induced neuropathy, fully preventing tactile hypersensitivity and preserving nerve endings. These effects persisted up to 14 days, indicating sustained action. High-dose Carba1 showed no detectable toxicity in mice, as confirmed by blood markers (fig. S5). While further studies are needed to establish its minimum effective and maximum tolerated doses, the results suggest Carba1 has favorable pharmacokinetic and pharmacodynamic properties for therapeutic use, with no observed toxicity.

NAMPT activation in naïve cells after 2 h resulted in decreased NAD^+^ and increased GTP levels, reflecting metabolic changes and feedback mechanisms to maintain NAD homeostasis. Similar studies with NAT showed significant metabolic variations at 1.5 hours that differed from those at 6 hours (*34*). Future research should focus on dynamic metabolic rewiring in naive cells, normal neurons, and PTX-stressed neurons to better understand pathways activated by PTX and Carba1’s NAMPT activation.

NAMPT overexpression is observed in various cancers, with certain types relying on NAMPT for NAD^+^ biosynthesis, making it a promising target for developing future anti-cancer inhibitors. (for recent reviews see Wei *et al.* (*48*), Wen *et al.* (*14*) and Velma *et al.* (*37*)). If Carba1 or its analogs are used to prevent CIPN in the future, it is essential to confirm that NAMPT stimulation does not promote tumor growth or compromise the anticancer efficacy of chemotherapy agents. In these experiments, a slight, non-significant increase in the GI_50_ of HeLa cells with Cis co-administered with Carba1 compared to Cis alone may reflect Carba1’s activation of NAMPT. Encouragingly, NMN-induced NAD^+^ elevation has shown neuroprotection against Cis-induced neurogenesis defects without affecting tumor growth or Cis antitumor efficacy (*49*). This study confirms that Carba1 does not affect the efficacy of therapeutic PTX doses and has no pro-tumor effects, establishing it as a strong candidate for CIPN prevention.

Among the NAMPT activators reported yet, (*14*), two classes of compounds, the P7C3 family with an aminopropyl carbazole core and the NAT family of hydroxyphenolamide derivatives, have been shown to prevent CIPN in animal models (*34*, *35*). While the crystal structure of the NAT-NAMPT complex has identified its binding site and activation mechanism (*34*), how P7C3 binds to and activates NAMPT remains unclear (*37*). We found that, like NAT, Carba1 could reverse cell death induced by the NAMPT inhibitor FK866, whereas P7C3 could not. Both Carba1 and NAT increased *in vitro* NAMPT activity, while P7C3 showed no detectable effect. These results align with previous studies that also failed to detect *in vitro* binding or activation of NAMPT by P7C3 (*50*–*52*). Although Carba1 shares a carbazole core with P7C3, it behaves more like NAT in its interaction with NAMPT. While we have shown that Carba1 binds directly to NAMPT through AS/MS, the crystal structure of the NAMPT-Carba1 complex is needed to fully understand the mechanism of NAMPT activation by Carba1.

Finally, we compared the different cellular activities of several Carba1 chemical analogs. Although some of these molecules may be differently metabolized by the cell, this comparative analysis is indicative of the structure-activity relationships of this carbazole series and relevant from a therapeutic point of view, since it is at the cellular level that drugs will ultimately act. The data suggest that compounds with either dual synergistic and neuroprotective activity or solely neuroprotective activity could be developed. However, the current data and available compounds do not allow us to predict the possibility of creating compounds with synergistic activity alone. Among the compounds studied, the one substituted on the nitrogen of the central ring (**16**) has a hybrid structure between Carba1 and P7C3 (fig. S3). Interestingly, this compound displayed none of the cellular activities analyzed, underscoring the differential nature of Carba1 and P7C3.

Despite these promising results, several limitations need to be taken into account when interpreting the findings and their wider implications. Firstly, preclinical studies have been conducted mainly *in vitro* and in animal models, which, while informative, may not fully replicate the complexity of human CIPN or cancer biology. The long-term safety of Carba1 remains uncertain and requires further evaluation in clinical settings. These limitations underline the need for further research to validate and extend these findings, particularly through rigorous preclinical and clinical studies.

Among the NAMPT activators described so far, only NAT has demonstrated neuroprotective activity. Carba1 therefore represent a new class of neuroprotective agents with this mechanism of action. Carba1 stands out as a groundbreaking therapeutic candidate, uniquely combining neuroprotective efficacy with synergistic activity alongside taxanes, addressing the critical unmet need for CIPN prevention without compromising anticancer efficacy. This dual action highlights Carba1’s potential to transform chemotherapy’s therapeutic landscape, offering hope if it reaches the clinic-after regulatory preclinical development and clinical trials- for improved quality of life and treatment outcomes for cancer patients.

## MATERIALS AND METHODS

### Study Design

This study aimed to evaluate the potential of Carba1, a novel sensitizing agent for paclitaxel (PTX), to enhance the efficacy of other chemotherapies, assess its neuroprotective effects against chemotherapy-induced peripheral neuropathy (CIPN), and elucidate its mechanism of action. Carba1’s synergy with various chemotherapeutic agents was tested *in vitro* using viability assays on HeLa cells, while its neuroprotective effects were evaluated *in vitro* on dissociated neurons and DRG explants from mice and rats, and *in vivo* using a rat model of PTX-induced neurotoxicity. The mechanism was investigated through untargeted metabolomics, NADH imaging, NAMPT activity assays, and AS-MS binding analysis. Its impact on tumor progression and PTX’s anti-tumor efficacy was assessed in a HeLa cell xenograft mouse model. All procedures adhered to ethical and regulatory standards, including NIH and FELASA guidelines, ARRIVE recommendations, and approvals from institutional and national committees (e.g., APAFIS#44532 and APAFIS#33137). Animal use was minimized, and no exclusions were made during analyses.

### Reagents

See Table S1.

### Animals

Primary cultures of dissociated adult DRG neurons were isolated from male and female C57Bl/6J mice, following protocols approved by Columbia University’s Institutional Animal Care and Use Committee in compliance with NIH guidelines. DRG explants were cultured from C57Bl/6J embryos (harvested at 13.5 days post-coitus) following FELASA-compliant procedures at the Institute for Advanced Biosciences. *In vivo* studies of CIPN were conducted on five-week-old Sprague Dawley rats (24 males and 24 females, Janvier Labs, France), housed in standard conditions (3 per cage) with *ad libitum* access to food and water under a 12:12 light/dark cycle. All experiments were conducted in accordance with the ARRIVE guideline (*53*). Ethical approval for these experiments was granted by the Auvergne Animal Care and Use Committee and the French Ministry of Higher Education and Research (APAFIS#44532), with efforts to minimize animal use. Tumor growth analyses were performed on 6-week-old female NMRI nude mice (Janvier Labs), housed at the Institute for Advanced Biosciences under ethical authorization from the Grenoble Ethics Committee (APAFIS#33137).

### Preparation and staining of dissociated neurons from adult DRGs

DRGs were dissected from 8- to 10-week-old mice in cold Hank’s balanced salt solution (HBSS) or Dulbecco’s Modified Eagle’s medium (DMEM) and dissociated in 1 mg/mL Collagenase P for 1 h at 37 °C, followed by 0.05% trypsin for 3 min at 37 °C. Dissociated cells were washed with Neurobasal medium supplemented with 2% B-27 Plus Supplement, 0.5 mM glutamine, 10% FBS, and 100 µg/mL penicillin-streptomycin. Neurons were then triturated by repeated gentle pipetting until there were no more visible clumps, resuspended in Neurobasal medium with 10% FBS and plated onto 12-well plates (over Ø18 mm coverslips) that have previously been coated overnight with 100 μg/mL poly-D-lysine at 37 °C and for 1 h at 37 °C with 10 μg/mL laminin. After 4 days *in vitro* (DIV), at least 30% of media was changed and 10 μM AraC (Cytosine β-D-arabino furanoside) was added to suppress non-neuronal growth. At DIV 12, neurons were treated for 72 h with 12 µM Carba1, alone or in combination with 50 nM PTX, 10 µM Cis, or 100 nM Bort. Neurons were fixed with 4% paraformaldehyde, washed with PBS, permeabilized with PBS-0.01% Triton X-100 for 15 min, and blocked with PBS-10% FBS for 1 h before being stained overnight at 4°C with neurofilament antibody. Following PBS washes, secondary antibody (Goat anti-Chicken AlexaFluor 488) was applied for 2 h at room temperature (R.T.). Following additional PBS washes, coverslips were mounted with Fluormount-G.

### Preparation and staining of embryonic 3D DRG explants

Embryonic DRG 3D explants were prepared from E13.5 mouse embryos and seeded at 1 explant/1.9 cm² on acid-treated, Matrigel-coated glass coverslips. Explants were cultured at 37°C under 5% CO₂ and high saturated in humidity. From DIV 0 to DIV 3, cultures were maintained in C-medium supplemented with 10% FBS, 1X GlutaMAX, 4 g/L D-glucose, 50 ng/mL 2.5S nerve growth factor (NGF) and 1% penicillin/streptomycin. From DIV 3 to DIV 7, neurobasal medium supplemented with 1X B27, 1X GlutaMAX, 4 g/L D-glucose, 50 ng/mL 2.5S NGF and 1% penicillin/streptomycin was used to promote axon development. From DIV 7 onward, differentiation medium (C-medium supplemented with 5 µM of forskolin, 50 µg/mL ascorbic acid and 25 µg/mL heparin) was used to support glial cell maturation and myelination, with media were changed every 2 days.

At DIV 11, explants were treated for 72h either with DMSO (control), 12 µM Carba1, 500 nM PTX, or a combination of both. DRGs were fixed with 10% formalin for 20 min at R.T., washed with PBS and permeabilized for 10 min with PBS-Triton 0.1%. Blocking was performed using a solution of 3% Bovine Serum Albumin, 0.1% glycine and 5% Goat Pre-Immune serum in PBS for 2h at R.T. DRGs were then washed in PBS and incubated overnight at 4°C with of anti-TUBB3 (1 µg/mL) and anti-MBP (1 µg/mL). After PBS washes, secondary antibodies (goat anti-mouse AlexaFluor 488, 1:1000, and goat anti-rabbit AlexaFluor 647, 1:1000) were applied for 1h at R.T. Explants were mounted with Prolong Gold antifade reagent with DAPI.

### Primary culture of rat sensory neurons (SN) and Schwann cells (SC)

These experiments, conducted by Neuro-Sys company (https://www.neuro-sys.com/) involved primary sensory neurons and Schwann cells cultured as previously described (*54*).

On DIV 19, cultures were pre-treated with Carba1 (1 to 10 µM) for 1 h before exposure to Cis (10 μg/mL) for 24 h in presence of Carba1. On DIV 20 the medium containing Cis was replaced with fresh medium and Carba1 for another 24 h, ending the culture on DIV 21. Cells were fixed with ethanol/acetic acid (95%/5%) for 5 min, permeabilized, and blocked with PBS containing 0.1% saponin and 10% FBS. Cells were incubated with monoclonal anti-MBP (1:1000) and polyclonal anti-neurofilament (1:500) antibodies for 2 h, followed by CF 488 A goat anti-mouse (1:400, Sigma) and AlexaFluor 568 goat anti-rabbit (1:400, Molecular Probe) for 1 h at R.T. Hoechst was used to stain nuclei.

Thirty images per well were acquired at 20X magnification using ImageXpress (Molecular Devices) with identical acquisition settings, and analysis was automated with MetaXpress® software. Key parameters measured included total neurite length (NF, μm), total neurons (NF^+^ cells), and myelination (MBP-NF overlap, μm²).

### Animal model of paclitaxel-induced peripheral neuropathy and Carba1 administration

This protocol, adapted from Balayssac *et al.* (*25*), involved intraperitoneal injections of PTX (5 mg/kg) on days 2, 4, 7, and 9 for a cumulative dose of 20 mg/kg (Fig. 2F). PTX was prepared by diluting a 20 mg/mL stock solution (cremophor®EL/ethanol, 1/1) to 5 mg/mL in 0.9% NaCl. Control animals received an equivalent volume of the vehicle (cremophor^®^EL / ethanol, 1/1) diluted 1/4 in 0.9% NaCl.

Carba1 was injected intraperitoneally 6 times (50 mg/kg) on days 0, 1, 2, 4, 7, and 9, resulting in a cumulative dose of 300 mg/kg (Fig. 2F). For each injection, Carba1 was injected intraperitoneally at 50 mg/kg on days 0, 1, 2, 4, 7, and 9, for a cumulative dose of 300 mg/kg, using a 150 mg/mL stock solution (cremophor®EL/ethanol, 1/1) diluted to 50 mg/mL in 0.9% NaCl. Control animals for Carba1 received the same volume of the vehicle (cremophor^®^EL/ethanol (1/1) diluted 1/3 in 0.9% NaCl). Four treatment groups were defined: control (Carba1 vehicle + PTX vehicle), Carba1 (Carba1 + PTX vehicle), PTX (Carba1 vehicle + PTX), and Carba1+PTX. To prevent cross-exposure through feces and urine, treatment groups were randomized by cage, with each cage housing 3 animals.

### Statistical analysis

Each *in vitro* experiment was performed at least 3 times independently. Data shown in each figure were analyzed by appropriate statistical tests specified in the figure legends.

## List of Supplementary Materials

Table S1

Fig S1 to S5

Supplementary legends

Supplementary methods

## Acknowledgments

We acknowledge the MicroCell facility (GIS IBiSA, ISdV, IAB), member of the national infrastructure France-BioImaging supported by the French National Research Agency (ANR-10-INBS-04)) for optical microscopy.

We are grateful to M.C. Laisne for her initial input in the project and to Dr Pascal Mossuz for his help in interpreting clinical data (blood counts, markers of renal or hepatic function) from the mouse xenograft tumor experiment.

## Funding

Société d’Accélération de Transfert de Technologies Linksium Maturation grant CM210005, (L.L).

Ligue contre le Cancer Allier et Isère (LL, C.T, D.B.)

Fondation ARC pour la recherche sur le cancer (CT)

Sanofi pilot grant (FB)

National Institutes of Health grant R01CA279401A1 (FB)

National Institutes of Health grant RF1AG050658 (FB)

French National Research Agency ANR-17-EURE-0003, doctoral fellowship (JA).

## Author contributions

Conceptualization: LL, FB, CT, DB

Formal analysis: BEH, JVi

Funding acquisition: LL, FB, CT

Investigation: LB, MEP, DB, NJ, JA, ST, CT, JVi, PS, PD, JVo, SMi, SMa, AM

Methodology: LB, MEP, UW, CC, PD, JA, DB, VJ

Project administration: LL, LB, FB, PD

Resources: PS, PD

Supervision: ST, CT

Validation: LB, LL, FB, DB

Vizualisation: LL, LB, BEH, PD

Writing, original draft: LL, FB, LB

Writing - review & editing: LL, FB, CT, PD, AM

## Data and materials availability

All data are available in the main text or the supplementary materials.

