## supplementary data for "Preventing neuropathy and improving anti-cancer chemotherapy with a carbazole-based compound"

### Legends of supplementary figures

**Fig. S1**

(**A**) Representative images of neuronal staining (TUBB3) in axons of 3D DRG mouse explants treated for 72 h with DMSO (Control), 500 nM of paclitaxel (PTX), 12 µM of Carba1 or their combination as indicated. Scale bar, 20 µm.

(**B**) Representative images of neuronal (TUBB3, green) and Myelin Basic Protein staining (MBP, red) from 3D DRG mouse explants treated for 72 h with DMSO (Control), 500 nM of PTX, 12 µM of Carba1or their combination as indicated. Scale bar, 100 µm.

(**C**) Effects of the different indicated treatments on weight gains of rats in a model of PTX induced neuropathy. Control animals (black curves) received vehicle injections. Weight of animals injected with 50 mg/kg Carba1 (blue curve), with 5 mg/kg PTX (red curve) and Carba1 together with PTX (orange curve), according to the experimental design at the indicated days is shown in Figure 2F. No significant difference (ANOVA) was observed.

**Fig. S2**

Validation of the OPLS-DA model of Fig 4A-B: permutation testing with 999 permutations.

**Fig. S3**

(**A**) Structure of Carba.

(**B**) Structure of P7C3. The common carbazole core is highlighted in orange.

**Fig. S4**

Weight of mice treated with PTX (8 mg/kg, red), Carba1 (60 mg/kg, blue), the combination (orange) of PTX (8 mg/kg) and Carba1 (60 mg/kg), or the vehicle (black). Dotted lines indicate treatment injection days.

**Fig. S5**

(**A**) Mouse blood cell counts

(**B**) Mouse renal parameters

(**C**) Mouse metabolic parameters

(**D**) Mouse liver parameter

### Supp Figure 1

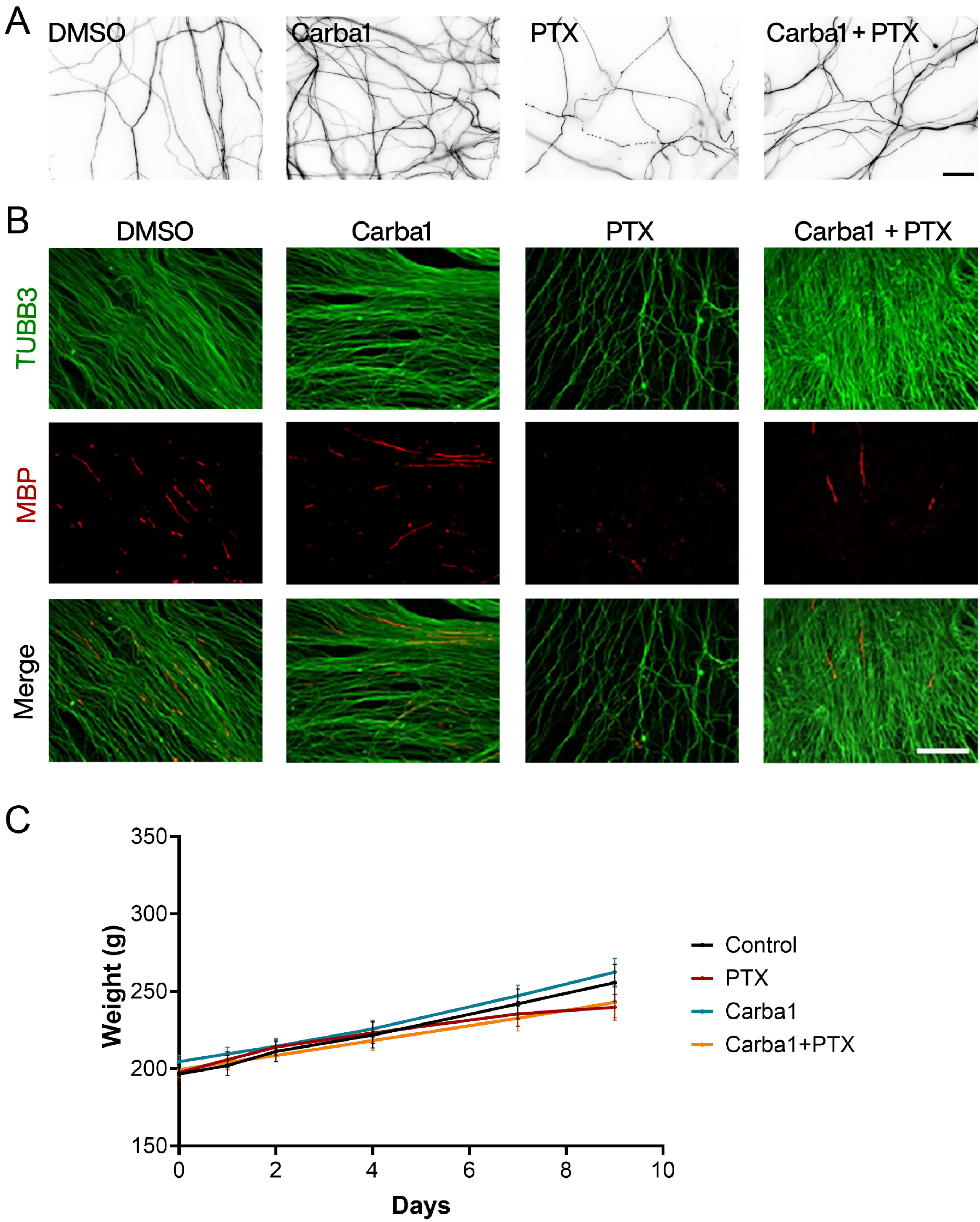

### Supp Figure 2

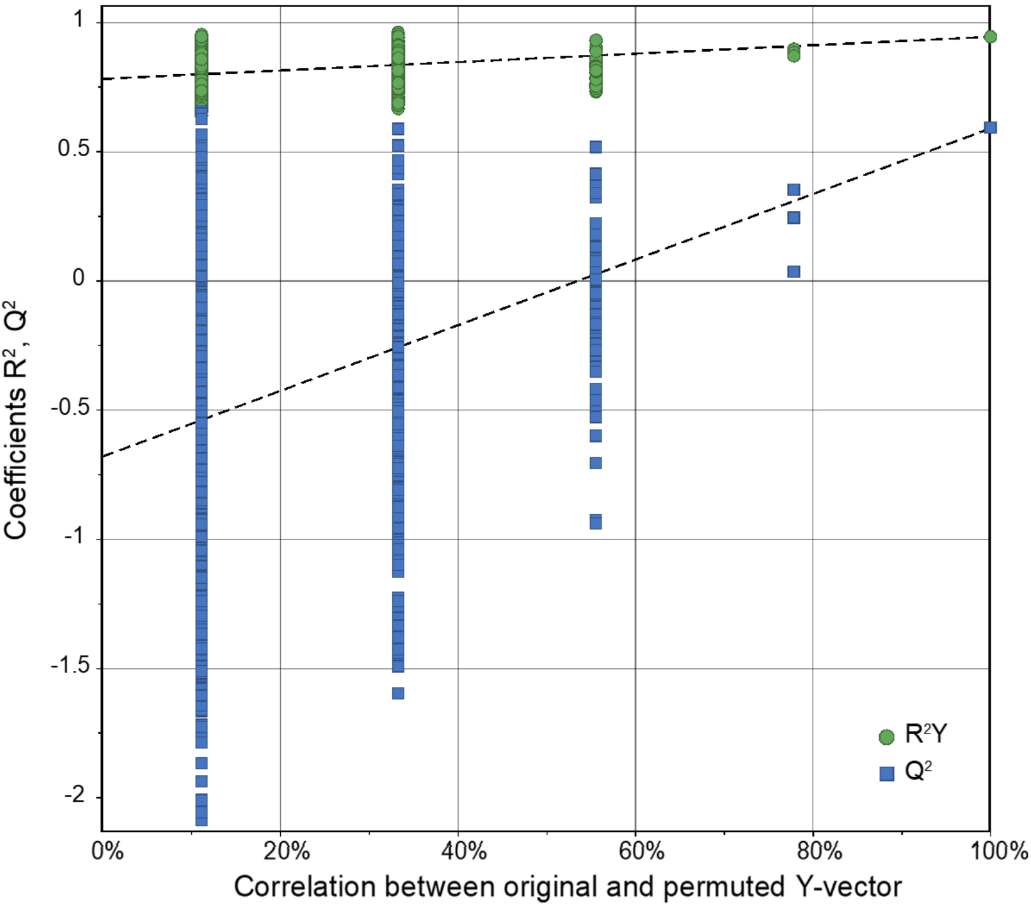

### Supp Figure 3

A Carba1

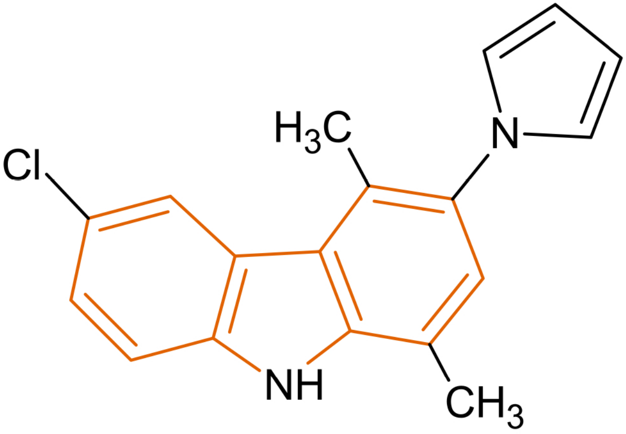


B P7C3

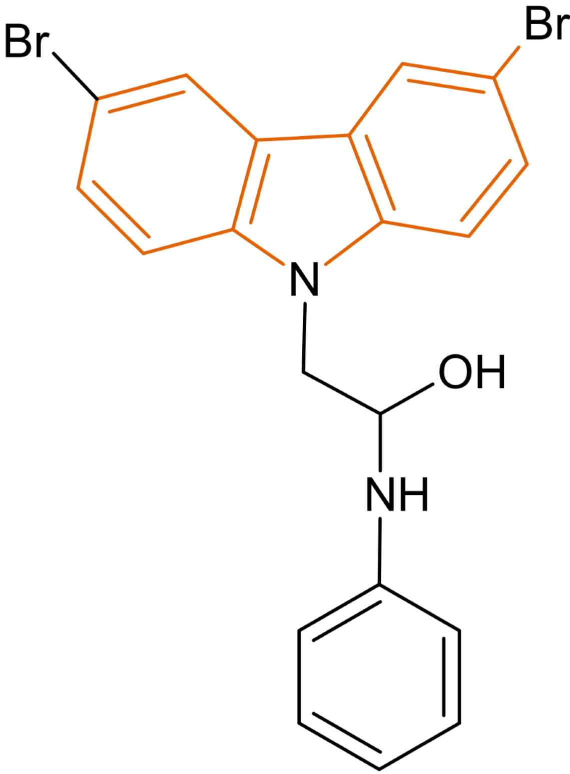

### Supp Figure 4

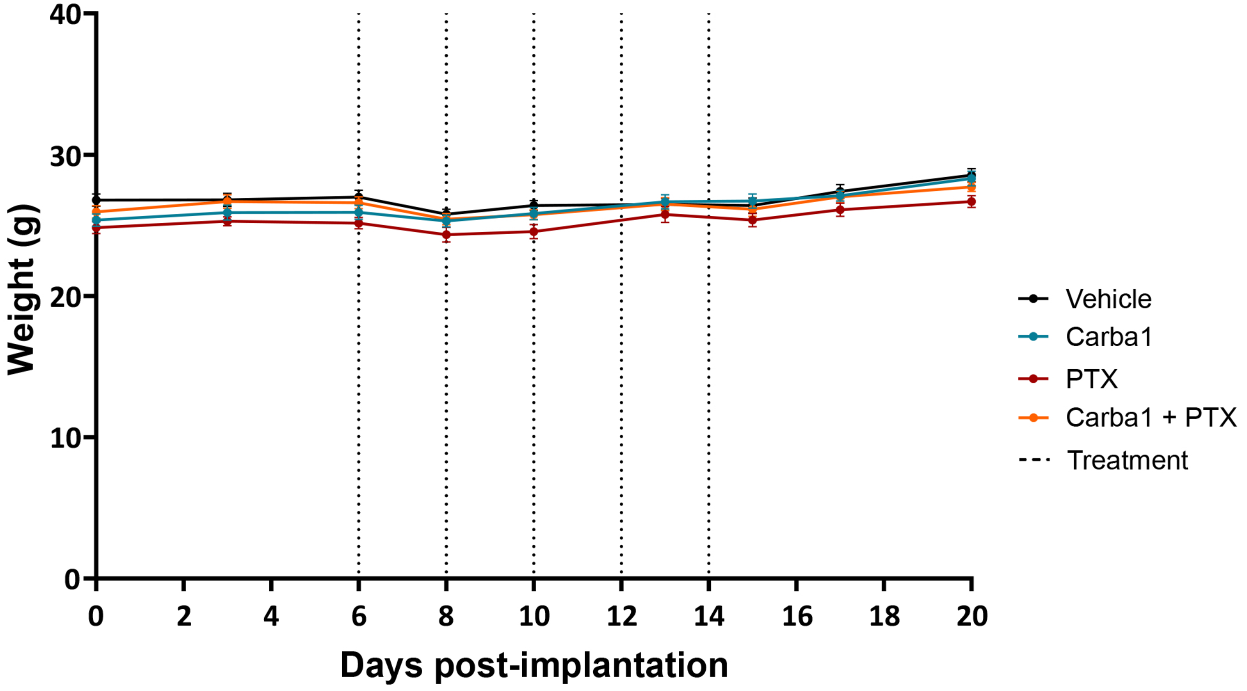

### Supp Figure 5

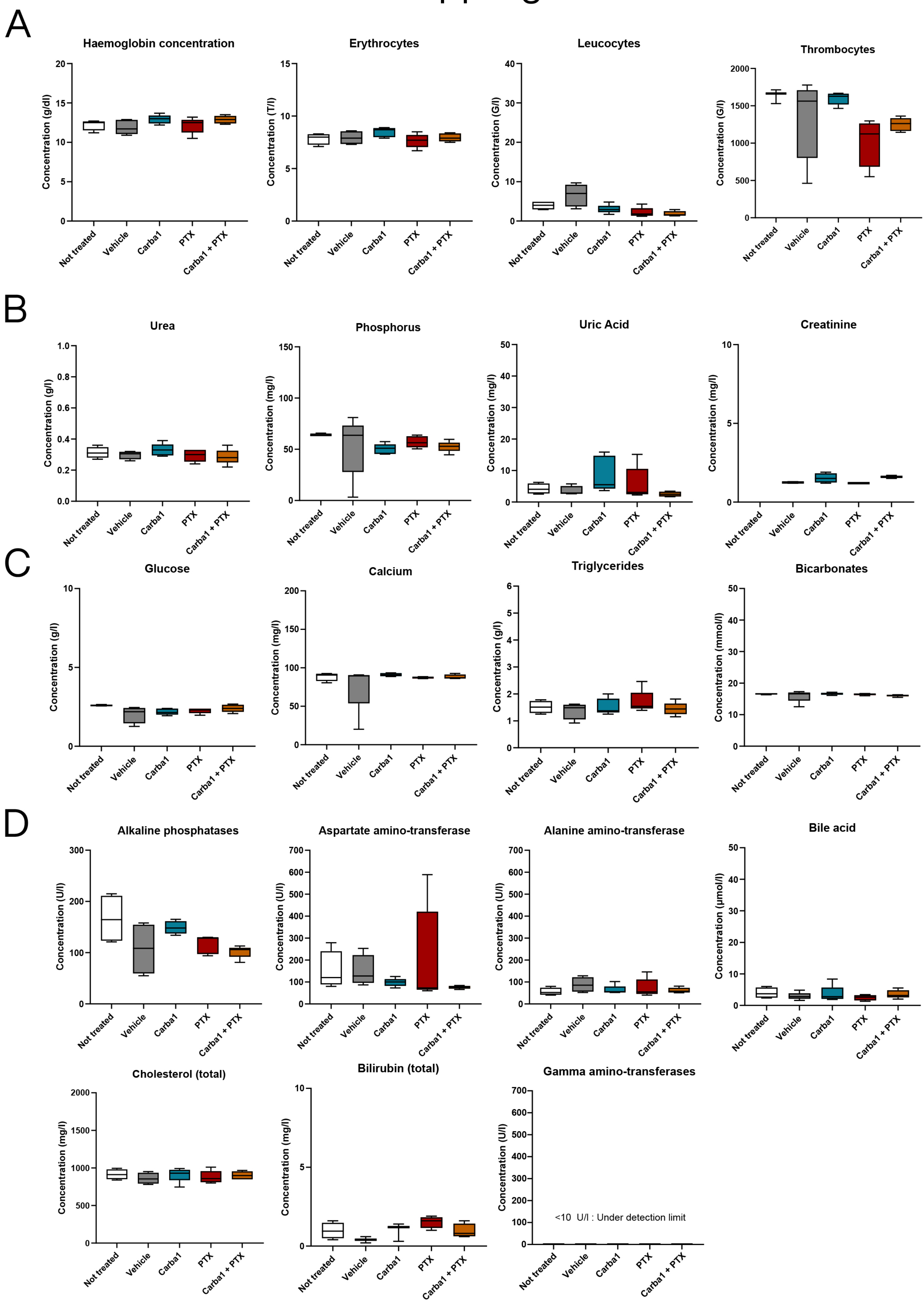

**Table S1 : list of reagents, materials and antibodies used in the study**

| Product | Supplier | Reference |
|---|---|---|
| 4-chamber Labtek | Dutscher | 55086 |
| 96-well microplates | Greiner | 655077 |
| Anhydrous dimethyl sulfoxide (DMSO) | Sigma-Aldrich | D4540 |
| AraC (Cytosine β-D-arabino furanoside) | Sigma-Aldrich | 147-94-4 |
| Ascorbic acid | Sigma-Aldrich | A92902 |
| B-27 Plus Supplement | Gibco | 175040441X |
| Bortezomib | Sigma-Aldrich | 5.04314 |
| BSA (Bovine Serum Albumin) | Sigma-Aldrich | A3912 |
| Carba1 | CERMN and Edelris | |
| Cisplatine (Cis) | Sigma-Aldrich | 232120 |
| Collagenase P | Sigma-Aldrich | 11249002001 |
| CremophorEL | Sigma-Aldrich | C5135 |
| D-glucose | Gibco | A2494001 |
| DMEM (no phenol red) | Gibco | 31053028 |
| Docetaxel (DTX) | Sigma-Aldrich | Y0001466 |
| Dulbecco's Modified Eagle's medium (DMEM) | Life Technologies | 1249015 |
| Epothilone-B (Epo-B) | Sigma-Aldrich | E2656 |
| Fœtal Bovine Serum | Dutscher | S1900-500C |
| FK866 | Merck Millipore | 481908 |
| Fluormount-G | Southern Biotech | 0100-01 |
| Formalin solution | Sigma-Aldrich | HT5012 |

| **Product** | **Supplier** | **Reference** |
|---|---|---|
| Forskolin | Sigma-Aldrich | 93049 |
| GlutaMAX™ Supplement | Invitrogen | 35050061 |
| Glycine | Euromedex | 26-128-6405-C |
| Goat Pre-Immune serum | Gibco | 16210-064 |
| Hank's balanced salt solution (HBSS) | Life Technologies | 14170112 |
| Heparin | Sigma-Aldrich | PHR8927 |
| Hoechst | Sigma-Aldrich | H33258 |
| Laminin | Life Technologies | L2020 |
| Matrigel Growth Factor Reduced | Corning | 356230 |
| MEM | Gibco | 11090081 |
| Nab-Paclitaxel | Gift from CHU Grenoble Alpes | |
| NAMPT Activity Assay | Abcam | ab221819 |
| NAT | MedChemExpress | HY-144778 |
| Neurobasal™ Medium | Gibco | 21103049 |
| Neurofilament 200kD | Aves Lab | NFH |
| NGF | Sigma-Aldrich | N6009 |
| P7C3 | Sigma-Aldrich | D8446 |
| Paclitaxel (PTX) | Sigma-Aldrich | T7402 |
| Paclitaxel (PTX, for *in vivo* rat experiments) | Leancare | |
| PBS | Gibco | 70011044 |
| Penicillin/Streptomycin | Gibco | 15140122 |
| PFA | Sigma-Aldrich | P6148 |

| Product | Supplier | Reference |
|---|---|---|
| poly-D-lysine | Sigma-Aldrich | P1149 |
| PrestoBlue | Invitrogen | A13262 |
| Prolong Gold antifade reagent with DAPI | Invitrogen | P36935 |
| RPMI 1640 | Gibco | 61870036 |
| Triton X-100 | Sigma-Aldrich | T8787 |
| Trypsin | Life Technologies | 25300054 |
| **Antibodies** | | |
| Anti-mouse AlexaFluor 488 | Invitrogen | A11029 |
| Anti-protein gene product 9.5 (PGP9.5, rabbit) | Abcam | ab15503 |
| Goat anti-chicken AlexaFluor 488 | ThermoFisher Scientific | A-11039 |
| Goat anti-rabbit AlexaFluor 647 | Invitrogen | A21245 |
| Goat anti-rabbit AlexaFluor 488 | Invitrogen | A11008 |
| Anti-TUBB3 (mouse) | Covance | MMS-435P |
| Anti-MBP (rabbit) | Abcam | ab40390 |

**Supplementary methods**

**Cell lines**

The human HeLa cell line was tested negative for mycoplasma contamination. HeLa cells were grown in RPMI 1640 medium supplemented with 100 µg/mL penicillin/streptomycin and 10% Fetal Bovine Serum, and maintained in a humidified incubator at 37°C and 5% CO2.

**Cell viability assay**

Cell viability was analyzed using the colorimetric Prestoblue assay. Cells were seeded in 96-well microplates at a density of 7,500 cells per well, allowed to adhere for 24 h and treated for 72 h with either DMSO or drugs at indicated concentrations. Prestoblue (10 µL) was then added to each well, followed by a 30-minute incubation. Fluorescence was measured using a FLUOstar Optima microplate reader (Excitation, 544 nm; Emission, 580 nm, BMG Labtech).

**Degeneration index measurements in DRG neurons and DRG 3D explants**

Images of random fields of dissociated adult DRG neurons, immunostained with anti-neurofilament or anti-TUBB3 antibodies were captured using a 20X objective lens (Olympus IX81) and a Sensicam QE monochrome camera (Cooke Corporation). Axonal degeneration was quantified by measuring the total axonal area and fragmented axonal area in the same field. Images were processed in ImageJ using global auto-thresholding, binarization, and the particle analyzer module to detect fragments (20 to 10,000 pixels), as previously described (*16*). The degeneration index was calculated as the ratio of fragmented axonal area to total axonal area.

**Quantification of myelin segments in DRGs explants**

Images of random fields of fixed DRG explants immunostained for TUBB3 and MBP were acquired with consistent exposure times across samples using a Zeiss AxioImager M2 microscope and a Hamamatsu Orca R2 camera. Analysis was conducted in Fiji software, where images were thresholded using the default global auto-threshold method. $MBP^+$ segments in each field were counted and normalized to the $TUBB3^+$ area.

**Assessment of nociceptive disorders (tactile allodynia)**

Tactile allodynia, a common sensory disorder reported in animal models of CIPN (*55*), was assessed using an electronic von Frey test (Bioseb) (*56*). Rats were placed in plastic compartments on a wire floor and allowed to habituate for 15 minutes. The von Frey apparatus, with a plastic tip connected to a force transducer, was applied perpendicularly to the right hind paw, and force was gradually increased until paw withdrawal. The maximum force (in grams) causing withdrawal was recorded automatically. The average of two measurements, not differing by more than 10 grams, was used as the nociceptive threshold (*56*). Rats were habituated to the compartments three days before the first injection, and the experimenter was blinded to treatment groups.

**Assessment of serum neurofilament light chain concentration**

Blood samples were collected on day 15 in vials with a clot activator, left at R.T. for up to 1 h before centrifugation at 4100 rpm at 20°C for 10 min. Serum was then collected, aliquoted, and stored at -80°C until analysis. To reduce batch effects, serum samples were randomized on assay plates. Neurofilament light chain (NfL) concentration was measured in the serum using commercial kits (Simple Plex Ella® ProteinSimple, USA; Rat NfL Kit ProteinSimple) (*25*).

**Assessment of intra-epidermal nerve fiber density**

Skin samples from both hind paws were fixed in 4% PFA overnight at 4°C, followed by sucrose cryoprotection (10%, 20%, and 30%) at 4°C overnight. After embedding in tissue freezing medium (TFM), the samples were at -80°C. Sections (20 µm) were cut using a cryostat at -17°C, washed with PBS for 5 min and dried. The sections were blocked in PBS/0.2%Triton / 1%BSA for 1 h, and then incubated overnight at room temperature with anti-PGP9.5 antibody (1/400). After washing with PBS, sections were incubated with goat anti-rabbit AlexaFluor488 secondary antibody for 2 h, and the nuclei were stained with DAPI. Imaging was performed by acquiring 0.5 µm Z-stacks and exposure times for DAPI and FITC set to 600 ms. Quantification followed the method described by Lauria et al. (*57*).

**NMR metabolomics analyses**

1x10$^6$ HeLa cells were seeded in Ø100 mm cell culture dishes and allowed to adhere for 48 h. Cells were then treated with DMSO or 12 µM Carba1 for 2 h at 37°C and 5% $CO_2$. After treatment, the medium was removed, and cells were washed with PBS at R.T. before quenching with methanol at -20°C and scrapping (*58*). Dried extracts were obtained by evaporating the solvent under gentle $N_2$ flow and stored at -80°C until analysis. For NMR, extracts were resuspended in 620 µL 100% $D_2O$ phosphate buffer (pH=7.4) and 550 µL was transferred into 5 mm NMR tubes. Untargeted NMR analysis was performed at 27°C on a Bruker Avance IVDr spectrometer (600 MHz, $^1$H resonance) with a conventional BBI probe. $^1$H metabolic profiles were acquired using a NOESY experiment with water presaturation (Bruker pulse program noesygppr1d) and a spectral width of 11904.792 Hz. Experimental parameters included a 10 ms mixing time, 2 s acquisition time, 3 s recycle delay, and 512 scans. The 90°s pulse length was calibrated at ~11.09 µs per sample after automatic shimming and tuning. NMR free induction decays were multiplied by an exponential function corresponding to a line broadening of 0.3 Hz prior to Fourier transform. Spectra were phased, baseline corrected, referenced to the alanine doublet at 1.47 ppm, and bucketed into 9900 variables (10$^{-3}$ ppm-wide) excluding water and methanol signals before multivariate statistical analysis. Data were normalized to the total sum of intensities. OPLS (orthogonal projection on latent structure) discriminant analysis was carried out on Pareto-scaled variables using SIMCA® 17 (Sartorius). Models were validated using 7-fold cross-validation and permutation testing. Individual metabolite levels were estimated by interactive line fitting using ChenomX NMR Suite 10 (ChenomX, Edmonton, Canada), and subsequent univariate statistics were obtained using GraphPad Prism.

**Imaging and quantification of endogenous NAD(P)H production**

HeLa cells were seeded in 4-chamber Labtek plates at 10,000 cells per well. After 48 h, cells were pre-incubated with or without 12 µM Carba1 for 2 h before addition of FK866 (5 and 10 nM). Drugs were added to phenol red-free DMEM medium, supplemented with 10% FBS 1% Penicillin/Streptomycin. After 24 h of drug incubation, cells were imaged using a confocal microscope (LSM 710, Zeiss) with a two-photon excitation at 700 nm (Chameleon Vision Laser, Coherent) and a 40x/1.2 W objective. Detection was performed in single-photon mode using an avalanche photodiode detector (ConfoCor 3). NAD(P)H signal intensity was quantified as the mean signal per cell using ImageJ and normalized to the cell area.

**NAMPT activity assay**

NAMPT activity was measured using the commercial kit NAMPT Activity Assay according to the manufacturer's instructions. In brief, NAMPT was incubated with various concentrations of the compounds in the presence of ATP, nicotinamide, nicotinamide mononucleotide adenylyltransferase 1 (NMNAT1), and phosphoribosyl pyrophosphate (PRPP) at 30°C for 15 min to allow the production of NAD. The reaction was stopped by adding 20 µM of FK866 (NAMPT inhibitor). A mixture of water-soluble tetrazolium salts (WST-1), alcohol dehydrogenase (ADH), diaphorase, and ethanol was added and the absorbance ($OD_{450}$) measured for 90 min using the microplate reader ClarioSTAR (BMG Labtech).

**Assay for NAMPT-compound binding**

Affinity selection-mass spectrometry (AS-MS) was used to assess compound binding to NAMPT. AS-MS was performed by Edelris SAS (https://www.edelris.com/), which has developed the commercial AS-MS service 'EDEN platform', based on Zehender & Mayr (*59*). Conditions consisted of protein (NAMPT and Carbonic Anhydrase) at 3 µM and compounds (Carba1 and FK866) at 10 µM dissolved in corporate buffer.

**Tumor xenograft in mice**

Five-week-old NMRI nude mice (Janvier Labs) were implanted with 10 x $10^6$ HeLa cells (200 µl in PBS) on the right flank. On day 6, the mean tumor volume was 120 ± 4 $mm^3$. Mice were then treated every two days (D6, D8, D10, D12, D14) with intravenous injections of 200 µL of Carba1, PTX, or Carba1+PTX or vehicle (14% DMSO, 14% Tween-80, 72% PBS). Behavior and signs of pain were monitored daily. Body weight and tumor volume were measured 2-3 times per week using electronic calipers, with tumor volume ($mm^3$) calculated as: length (mm)/2 x $width^2$ ($mm^2$). Tumor growth inhibition was analyzed using GraphPad Prism with one-way ANOVA and Dunn's multiple comparison test for treatment *vs* vehicle. Mice were sacrificed on day 20, when the vehicle group reached ethical limits. Blood samples were collected by intracardiac puncture in EDTA coated tubes for differential blood counts and hemograms, outsourced to IDEXX BioAnalytics. For biochemical analysis, blood was collected in heparin-lithium tubes, and plasma was isolated by centrifugation and frozen at -80°C. Plasma was analyzed using MS Scan II (Melet Schloesing) and VET16 reagent rotors to measure markers of renal and hepatic function, metabolism, and nutritional and muscular status. Differential counts, hemograms, and biochemical data were analyzed with GraphPad Prism using Kruskal-Wallis test and Dunn's multiple comparison test.
